# Exploring the visual system of the Black Grouse (*Lyrurus tetrix*): Combining experimental and molecular approaches to inform strategies for reducing collisions

**DOI:** 10.1101/2025.01.12.632585

**Authors:** Simon Potier, Jean-Marc Lassance, Constance Blary, Justine Coulombier, Sandrine Berthillot, Jérôme Cavailhes, Charleyne Buisson, Virginie M. Dos Santos, Christine Andraud, Marjorie A. Liénard

**Affiliations:** Simon Potier, Expert Scientifique, Foucrainville, France; GIGA Institute, University of Liège, Belgium; CEFE, Univ Montpellier, CNRS, EPHE, IRD, Montpellier, France; Parc national de la Vanoise, Chambéry, France; Association Observatoire des Galliformes de Montagne, Sevrier, France; Centre de Recherche sur la Conservation des Collections, CNRS, Muséum National d’Histoire Naturelle, France; Laboratory of Molecular Biology of Sensory Systems, GIGA Institute, University of Liège, Belgium

**Keywords:** visual field, contrast vision, spectral sensitivity, opsin, bird collision

## Abstract

Understanding the visual systems of birds can inform conservation efforts and mitigate the impact of collisions with man-made structures. Here we investigate the visual abilities of the black grouse *Lyrurus tetrix* (Galliformes: Phasianidae), a European mountain bird highly vulnerable to collisions with aerial infrastructures. Using behavioural assays, we show that black grouse have wide monocular lateral visual fields, extensive binocular overlap and a minimal blind area behind the head, altogether indicating the ability to detect aerial objects from different fields of view. The spatial resolution and contrast sensitivity of the black grouse are in the low avian range but align with the ecology of a prey species and allow to establish the limits of detection of an object size and contrast under variable environmental conditions. To characterize the range of wavelengths perceived, the four cone visual pigments (SWS1, SWS2, Rh2, and LW) were reconstituted and characterized functionally, showing detection of light across a spectral range spanning the visible spectrum up to 650 nm and with limited sensitivity below 400 nm. Combined with spectral and achromatic modelling analyses, our results inform on the limits of detection of aerial objects and the perception of existing visual markers, offering practical perspectives to continue mitigating black grouse collisions.

## Introduction

Collisions are a recognised major source of bird mortality worldwide (Rosenberg et al., 2019, Loss et al., 2014), especially with glass or other reflective building surfaces (Machtans et al., 2013), wind turbines (Marques et al., 2014, Bernardino et al., 2018, Drewitt & Langston, 2008), power lines and aerial cables that intrude into the open airspace. With the rise of ski resorts in alpine regions, the development of road networks, ground and aerial infrastructures and cable transport systems (e.g., catex, cable cars, surface lifts) have substantially reshaped landscapes over the last fifty years, notably increasing habitat fragmentation and reducing natural wildlife habitats and refuges for mountain avifauna, in particular for ground-nesting birds (Watson & Moss, 2004, Segelbacher et al., 2008, Bech et al., 2013, Matysek et al., 2021). Ski resort areas have also been associated with high collision rates, especially for Galliformes (Miquet, 1990, Bevanger, 1998, Buffet & Dumont-Dayot, 2013).

The black grouse, *Lyrurus tetrix* (Phasianidae, previously *Tetrao tetrix*) is a highly sedentary ground-nesting gallinaceous bird that forages for terrestrial invertebrates, berries, plant shoots or leaves (Hambálková et al., 2024). The black grouse is sensitive to habitat change and human disturbances (Bernardino et al., 2018), and commonly exposed to a range of collision hazards. Documented effects include for instance deer fences in Scotland populations (Baines & Summers, 1997), power lines in Scandinavia (Bevanger, 1995), and ski lift cables in Alpine regions (Miquet, 1990, Buffet & Dumont-Dayot, 2013). When comparing black grouse population abundance in natural habitats versus ski resorts in the south-western Swiss alps, Patthey et al. (2008) quantified that habitat topology and ski lift density were already associated with 15% reduced local black grouse abundance but without linking it to collisions *per se* (Patthey et al., 2008). In the French alps, where the black grouse population is smaller and more isolated than in Scandinavian regions, around 400 deaths of black grouse have been recorded near aerial infrastructures since 1964, which represented ca. 70% of the total Galliformes death counts over that period (Corona & Dos Santos, 2023, OGM, 2010). According to earlier estimates of the French Alpine black grouse population in 2010 (OGM, 2010), collisions caused mortality in 2.5% of the population. This number is likely conservative since it excluded individuals that collide in flight but might die outside search areas, and since the probability of detecting carcasses is very low (Bech et al., 2012).

For more than 20 years, numerous ski resorts have equipped hazardous ski lifts with visual markers, small plastic or metallic devices (Figure 1, Table S1) designed to increase visibility through size, movement in the wind, reflective properties, and contrasting colour patterns (Dos Santos et al., 2021). However, the effectiveness of these markers can differ among target species and environmental conditions (Ferrer et al., 2020), and it remains unclear whether black grouse perceive hazardous obstacles introduced into their flight paths. To assess current solutions and possibly guide the development and efficient placement of visual markers, knowledge of avian species-specific vision, ecology and behaviour should necessarily be considered (Martin, 2011a, Martin, 2022, Martin & Banks, 2023).

**Figure 1.**
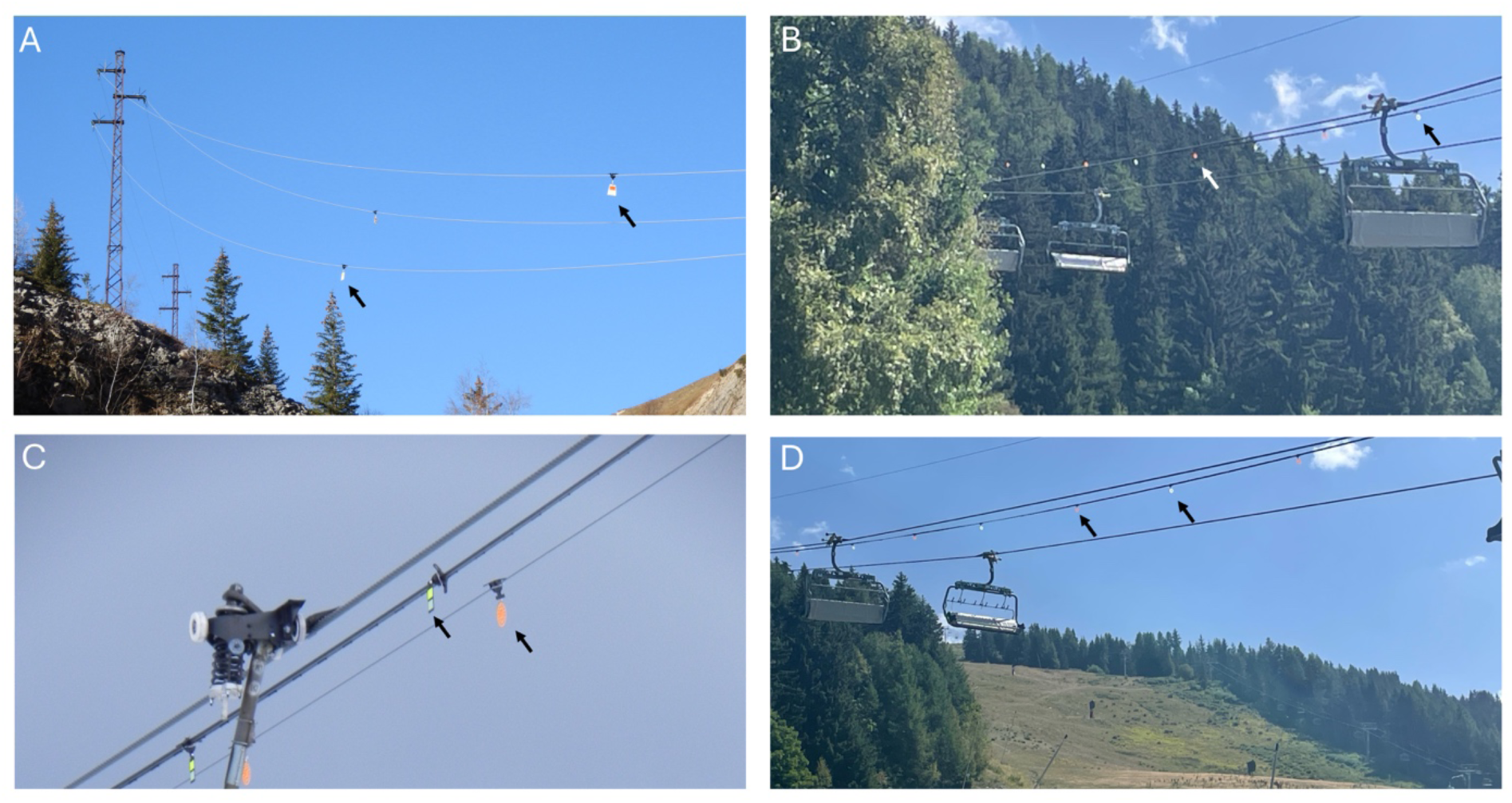
Aerial cables and ski lifts equipped with visual markers in the French Alps. Visual markers are indicated with arrows (see also Table S3 and Figure 7 for alternative designs). (A) Example of *black Firefly* markers (15 x 9 cm) on aerial electrical cables. (B-D) Examples of *Crocfast* and *Birdmarks* markers (13.5 cm in diameter) seen against (B) a green background, (C) a cloudy background, and (D) a clear sky. Photographs credit: VPN: A,C; JML: B, D.

Vision is the primary sensory modality for most avian species, collecting accurate and instantaneous environmental information from different directions and over varying distances (Land & Nilsson, 2012). Avian visual systems influence object detection for various tasks, and are shaped by natural history and ecology such as diel activity patterns, navigation, foraging strategies, or predator-prey interactions (Odeen & Håstad, 2010, Wu et al., 2016, Outomuro et al., 2017, Hauser & Chang, 2017, Cooney et al., 2019, Caves et al., 2024, Blary et al., 2024).

While object detection is influenced by its properties, the bird flight behaviour and the environmental light conditions, together with the bird’s visual characteristics appear to be important factors, including the fields of view, spatial resolution, as well as both its contrast and spectral sensitivity (Martin, 2022). The visual field is the volume of space around the head from which visual input is gathered at any given time (Martin, 2017a). It therefore determines whether an artificial structure or object falls within a blind area or within a bird’s field of view. In the latter case, the binocular region allows estimation of time before contact (Martin, 2009, Martin, 2017b, Martin et al., 2012).

An object must also reach a minimum size at a distance threshold to be detected prior to collision (Martin, 2022), which can be informed by determining spatial resolution. Spatial resolution is the ability to resolve two points as separate, and is a measure also used to quantify how well birds can distinguish fine details, defining their visual acuity (Land & Nilsson, 2012). Target detection further requires the ability to discern contrasts between an object and the background, or patterns within an object, and is defined by the eye contrast sensitivity. Finally, the spectral sensitivity of the avian eye is determined by different cone photoreceptors, which shape how birds detect wavelengths of light and perceive colours (Potier et al., 2020, Kelber, 2019, Stavenga & Wilts, 2014, Wu et al., 2016). Most avian visual systems have four single cone types that each contain one opsin visual pigment exhibiting distinct maximal wavelength sensitivity (λₘₐₓ) when forming complexes bound to 11,*cis*-retinal: SWS1 360-420 nm, SWS2 430-460 nm, Rh2 470-510 nm, LWS 530-560 nm (reviewed in Hagen et al., 2023). Bird cone opsins typically mediate a tetrachromatic colour vision system, in combination with cone-specific oil-droplet filters (Kelber, 2019). In birds, the short-wavelength sensitive 1 opsin (SWS1) absorbance range also varies from ultraviolet-sensitive to violet-sensitive (reviewed in Hauser et al., 2014, Hagen et al., 2023, Carvalho et al., 2007). Studying species-specific visual characteristics can inform about the limits of object detection in natural and human-made environments.

Despite the long interest in black grouse conservation efforts, we still know relatively little about its visual system. A behavioural study has shown that black grouse display a preference to eat naturally UV-reflecting waxy European blueberries (*Vaccinium myrtillus*) (Siitari et al., 2002). Physiological and molecular studies in other galliform species including chicken, quail, turkey and peacock also suggest a potential for some UV sensitivity in these species (Bowmaker et al., 1997, Hart et al., 1999, Cuthill et al., 2000). Despite these reports, our understanding of black grouse vision remains fragmentary, with many aspects of their visual capabilities still uncharacterised.

The aim of the present study was thus to quantify components of the black grouse visual system that impact object detectability. Specifically, we used a combination of reflex-like behaviours, ophthalmoscopy and molecular biology to assess the black grouse visual fields, spatial resolution, contrast sensitivity, and wavelength absorbance of cone visual pigments. We then measured the reflectance properties of anti-collision visual markers placed on ski lifts and discuss the black grouse visual system characteristics in the context of marker detection, as well as the risk of flight collision under variable visual environmental conditions.

## Material and methods

### (a) Birds and study location

Two 4-month-old male black grouse individuals belonging to the breeding centre of Frank Grosemans (Belgium) were used to study visual fields, spatial resolution and contrast sensitivity.

### (b) Visual fields

A non-invasive procedure was used to measure visual field characteristics in alert birds (Martin & Portugal, 2011). Each bird was held firmly for 20-30 min in a plastic restraining tube of the appropriate size to avoid any movement, with its legs cushioned by foam rubber and gently immobilized together with surgical tape (Micropore 1530/1B). The head was held following a natural position and placed at the centre of a visual field apparatus, a device that permits the eyes to be examined from known positions around the head. The bill was immobilized in a specifically manufactured steel and aluminium bill holder, and held with micropore tape, keeping the nostrils clear for normal breathing. The bill holder surfaces were coated with cured silicone sealant to provide a cushioned, non-slip surface. The bird’s head and eyes were photographed (iphone 13, Apple) while in the apparatus to determine the eye positions in the skull, the horizontal separation between the centre of both eyes, the eye-to-bill-tip distance and the bill length.

Visual field parameters were measured using an ophthalmoscopic reflex technique, with the perimeter’s coordinate system following latitude-longitude conventions and the equator aligned to the median sagittal plane, which divides the head vertically and symmetrically into left and right hemispheres. This coordinate system is used throughout the presentation of results. We first examined the eyes using an ophthalmoscope mounted on the perimeter arm with an accuracy of ± 0.5° to measure the boundaries of the retina projection from the positions that the eyes spontaneously adopted when they were fully rotated forwards, *i.e*. converged for estimation of binocular area boundaries, and backwards *i.e.* diverged for estimations of the blind area behind the head. The degree of eye movements and pecten projection were not measured to minimise restraint time.

We corrected our measurements to a hypothetical viewing point placed at infinity, based on the distance used in the visual field apparatus and the horizontal separation of the eyes (Martin, 1984). Using these corrected values, we constructed a topographical map of the visual field, including the lateral fields, binocular field, cyclopean field *i.e.* the total field around the head comprising the combined monocular fields of both eyes, and blind areas above and behind the head. The limits of the visual field were determined at 10° elevation intervals in an arc starting behind the head then above and in front of the head, down to an elevation 50° below the horizontal plane of the bill-tip. At elevations where the bill holder blocked our view, binocular field width was interpolated as the mean of the binocular field widths immediately above and below these elevations (Martin & Portugal, 2011).

### (c) Spatial resolution

Spatial resolution (SR), or visual acuity, is the ability to perceive spatial detail. SR can be estimated from retinal photoreceptor or ganglion cell density in the retina (Coimbra et al., 2012, Lisney et al., 2012b), estimates that closely correlate with behavioural measures of visual acuity across vertebrates (Caves et al., 2024). There is also a strong positive correlation between visual acuity and eye size, which is obtained by measuring corneal diameter. Corneal measures thus offer the advantage of being non-destructive, while providing estimates of visual acuity that are comparable to those obtained from behavioural estimation (Potier et al., 2016, Caves et al., 2024).

Corneal diameters (CD) were measured from calibrated eye photographs with ImageJ 1.41 (Schneider et al., 2012) to obtain Axial Length (AL, in mm) using the formula for diurnal animals (Hall & Ross, 2007):

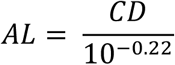

Spatial resolution (SR, in cycles per degree, abbreviated cpd) was calculated following the function derived from Caves et al. (2024):

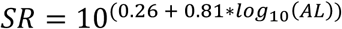

### (d) Contrast sensitivity

#### (i) Experimental set-up

We used an eye reflex-like behaviour setup to study the contrast sensitivity of each individual, following Wagner et al. (2022) and Blary et al. (2024). This approach yields peak contrast sensitivities comparable to those obtained with the operant conditioning method (Blary et al., 2024). The eye reflexes, or optocollic reflexes, are caused by involuntary compensatory mechanisms for image stabilization, *i.e.,* the bird follows the moving object by eyes to keep the image stationary on the retina (Wagner et al., 2022). Because most birds have limited eye movements between 2 and 18° (Martin, 2009, O’Rourke et al., 2010), they follow the object with large head movements. To estimate the optocollic reflex (OCR), birds were placed into a U-shaped box adapted to their size (60 x 40 x 44 cm; L x W x H), with a black wooden floor and ceiling, and clear polycarbonate walls (3 mm) (Figure 3A). The walls allow the bird to see the stimulus without influencing visual abilities (Blary et al., 2024). The ceiling consisted of two wooden boards spaced 5 cm apart: a perforated panel to allow air flow through and a smaller solid panel preventing the bird from seeing through the holes. All wooden surfaces were painted matte black (Liberty black Matt, GoodHome). A rear wooden door allowed the bird to be positioned inside the box. The box was placed at the centre of an octagon (radius 69cm) formed by computer monitors (Acer KG1 Series, 28 inches, 4K 3840x2160 px, 60 Hz) in portrait orientation, with their centres aligned to the level of the bird’s head. The stimulus covered 236 x 46°of the visual field (horizontal x vertical), sufficient to elicit an OCR (Schmid & Wildsoet, 1998, Blary et al., 2024).

The monitors and the bird were enclosed in an opaque black cloth tent (300 x 300 x 250 cm) with the monitor bases and floor also covered in black cloth to minimize external visual cues. The experimenter remained outside the tent throughout the experiment. A video camera (25 frames per second; Sony DCR-SR55 HD) positioned above the monitors recorded the bird’s behaviour and a smartphone (iphone 13, Apple) connected to an external screen allowed continuous remote monitoring.

#### (ii) Stimuli

Stimuli consisted of vertical sinusoidal achromatic gratings of varying spatial frequencies and contrasts generated in Matlab (R2021) using Psychtoolbox (Brainard, 1997). Monitors were driven by MacBook Pro (Apple) via an 8-Port 4K/60Hz HDMI splitter (ST128HD20, StarTech). The angular velocity of the gratings was set to 20° s^-1^, matching values used in closely related species (Blary et al., 2024). To obtain the contrast sensitivity function (CSF) – including peak contrast sensitivity, its corresponding spatial frequency, and the high cut-off frequency (*i.e.* the highest spatial frequency detected at a Michelson contrast of 1) – we used six spatial frequencies (0.47, 0.56, 0.85, 1.06, 1.27 and 1.59 cpd) and Michelson contrasts ranging from 0.012 and 0.99 (Table S2). Specifically, Michelson contrast is defined as:

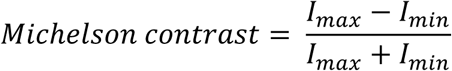

where *I_min_* and *I_max_* are the minimum and maximum luminance; values range from 0% (no contrast) to 100% (perfect black-white difference). For each contrast level, luminance differences between the darkest and lightest regions of the monitor centre were measured with a luxmeter (Hagner ScreenMaster, B. Hagner, Sweden) and expressed in candelas per square meter (cd/m^2^).

Each stimulus was displayed for 20 s (Schmid & Wildsoet, 1998, Shi & Stell, 2013, Wagner et al., 2022, Blary et al., 2024), with stimulus direction (clockwise or anticlockwise) varied in a pseudorandom order, limiting consecutive identical directions to three. Illuminance at the bird’s head was 110 lux inside the U-shaped box and 321 lux outside, and stimulus brightness was 73 cd/m^2^.

#### (iii) Experimental procedure

Computer monitors were switched on 20 minutes before to the experiment to reach maximum brightness. Following the bird’s placement in the device, a 10-minute habituation period was observed before the experimental session began. The experimenter visually assessed the presence of the optocollic reflex (OCR) in accordance with standard practice (Schmid & Wildsoet, 1998, Shi & Stell, 2013, Blary et al., 2024). Each session began with maximum contrast (Michelson contrast = 0.99) to ensure that the bird was expressing an OCR detectable by the experimenter. Contrast was then varied stepwise as described in Blary et al. (2024), to identify the threshold at which birds no longer exhibited the reflex. Each trial combined a specific combination of spatial frequency and contrast and was separated by at least 5 seconds of an isoluminant homogeneous grey screen (longer if the bird did not look at the monitors). We applied the “four of five” criterion: each combination was presented five times, and a response was considered reliable if the OCR occurred in at least 4 trials (Shi & Stell, 2013, Blary et al., 2024). The lowest contrast meeting this criterion was defined as the threshold contrast. Once determined, the experiment was repeated at the same angular velocity with a different spatial frequency. All session were video-recorded, allowing retrospective verification of OCR presence or absence.

### (e) Detection distance

Under optimal conditions, the distance at which an object can be detected and avoided depends on three factors: the spatial resolution, reaction time (t_react_) and flight speed (*v)* (Martin, 2022). In birds, it would seem reasonable to consider that a minimum t_react_ = 2s is necessary to initiate a change in flight path (Martin, 2022). Using the allometric relationship between body mass and cruising flight speed presented by Alerstam et al. (2007), the mean body mass of black grouse (1,068.7 g; Dunning 2007) predicts a typical *v* of 16.04 m·s⁻¹. If the object only needs to be visible during reaction, the minimum detection distance (D) of an object while in flight is D = 2*v*. Therefore, the minimal detection distance of an object is 2*16.04 = 32 m.

From this minimal detection distance, the spacing (d) between two objects (*e.g.* two adjacent visual markers) while actively flying can be estimated using the trigonometric relationship between angle and detection distance as follow: d = 2*D. tan(θ/2), where θ is the binocular overlap angle in radians.

Under optimal conditions, the minimum width of an object (*w*) to meet the spatial-frequency limit of one cycle can be estimated using 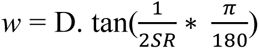, where *SR* = Spatial Resolution. A factor of 2 is applied to *SR* because visual acuity is expressed in cpd, and one cycle corresponds to two resolvable bars; birds can therefore discriminate two objects within a single cycle.

### (f) Opsin λmax characterization via heterologous expression

#### (i) Collection of eye tissues and extraction of ribonucleic acid (RNA)

Ocular tissue, including the retina, was dissected by a veterinarian from an accidentally killed adult female black grouse collected in Bourg-Saint-Maurice France (ID: JB-2021-E001, POIA Birdski Project 2020-2023 Vanoise), respecting permits in place. The collected tissue was directly preserved in DNA/RNA reagent (Zymo Research) and stored at -20° C. Before extraction, the tissue sample was removed from the stabilization solution, briefly dried and then ground to a fine powder in a mortar cooled with liquid nitrogen. We then proceeded with RNA extraction with a Monarch Total RNA miniprep kit (NEB) following the manufacturer’s instructions. The integrity of the RNA was evaluated using a bioanalyzer and the quantity assessed using a Qubit High Sensitivity RNA assay (Invitrogen).

#### (ii) cDNA libraries and RNA-sequencing

Five hundred micrograms of high-quality RNA were used to prepare a deoxyribonucleic acid (DNA) library following the protocol detailed in NEBNext Ultra II Directional RNA library prep kit for Illumina (NEB), including the following specifications: selection of messenger RNA (mRNA), a selected average size of 500 base pairs for fragmentation, and the addition of Illumina adapters. The quality of the cDNA library was evaluated using a bioanalyzer as well as by quantitative PCR (Polymerase Chain Reaction). Paired-end sequencing (2x150bp) was performed on an Illumina NovaSeq6000 equipped with an S4 flowcell (NovaSeq Control Software 1.8.0/RTA v3.4.4, SciLifeLab, Stockholm, Sweden). In total, we obtained 197.8 million read pairs. The transcriptome sequencing data have been deposited into the European Nucleotide Archive (ENA) under the project accession PRJEB86944.

#### (iii) Pre-assembly filtering

Raw reads were first preprocessed by correcting and excluding aberrant reads using the k-mer-based read-error correction tool Rcorrector (Song & Florea, 2015). We next proceeded to adapter removal and quality trimming (--quality-cutoff=20) using cutadapt version 4.2 (Martin, 2011b). During that step, we also removed read pairs with either read shorter than 36 bp. Next, we removed reads corresponding to rRNA sequences by mapping the reads against a custom database built from SILVA SSU and LSU datasets (release 138.1) (Quast et al., 2013) using bowtie2 (Langmead & Salzberg, 2012). Finally, reads corresponding to mitochondrial sequence were extracted using MITGARD (Nachtigall et al., 2021). The final set of assembly-ready reads contained 173.8 million read pairs.

#### (iv) *De novo* transcriptome assembly and annotation

De novo assembly of the pre-processed paired-end reads was performed using Trinity v2.14.0 (Garber et al., 2011) with default parameters. Completeness of the assembly was assessed with BUSCO v3.0.2 (Simão et al., 2015) using the Vertebrata and Aves datasets. We applied the pipeline implemented in Trinotate v3.2.1 (Bryant et al., 2017) to predict candidate protein coding regions within transcript sequences using TransDecoder v5.5.0 (https://github.com/TransDecoder/TransDecoder) and to generate a functional annotation of the transcriptome data. We retrieved full-length candidate opsin transcripts from the resulting annotation table (Dataset S1).

#### (vi) Opsin functional expression

Opsin-specific oligonucleotide primers (Table S3) encompassing two distinct restriction sites were designed to amplify the coding region of each identified visual opsin (SWS1, SWS2, Rh2, LW) by PCR using retinal cDNA synthesized using the GoScript protocol (Promega) as template. After verification by agarose gel electrophoresis, expected amplified product sizes were gel-purified using the Monarch DNA gel extraction kit (NEB), digested with restriction enzymes, subcloned into the linearized pcDNA5-FLAG-T2A-mruby2 expression vector (Liénard et al., 2021), and propagated into a Q5 strain of *Escherichia coli* bacteria (NEB). After incubation of individual bacterial colonies in Luria-Broth culture medium containing ampicillin, DNA plasmids were purified, and each construct sequence was verified by Sanger sequencing. A stock of 480 micrograms for each final plasmid was prepared following the ZymoPURE II Plasmid MidiPrep kit protocol (Zymo).

#### (vii) Protein purification, absorbance measurement and UV-VIS analyses

Each plasmid was transfected into human embryonic kidney cells (HEK293T, ThermoFisher) following the PASHE procedure (Liénard et al., 2022). Briefly, 11-*cis*-retinal was added in solution in the dark (under red light) 6h post transfection. Cells were collected and the total membrane fraction solubilized using 1% n-Dodecyl-D-Maltoside detergent (DDM) 48h after transfection, and the opsin fraction bound to FLAG resin and incubated overnight. The next day, the opsin-FLAG complex was purified and eluted using FLAG peptide, subsequently concentrated by centrifugation at 4000 rpm and 4°C for 50 min prior to ultraviolet-visible (UV-VIS) spectroscopy. The absorbance of each purified rhodopsin (opsin in complex with the chromophore) was measured in the dark from 1.5 μL aliquots using a NanoDrop 2000/2000c UV-VIS spectrophotometer (ThermoFisher). For each rhodopsin, we estimated lambda max by nonlinear fitting of the absorbance data using a visual pigment template (Govardovskii et al., 2000a). We performed 1,000 bootstrap replicates to compute lambda max estimates and the associated confidence intervals (Liénard et al., 2022) in R v. 3.6.6. using the rsample (Frick et al., 2022) and tidymodels (Kuhn & Wickham, 2020) packages.

### (f) Modeling of spectral sensitivity in cone photoreceptors

To derive a theoretical effective eye spectral sensitivity in the black grouse, which depends on the visual pigment opsin λ_max_ and oil droplet filtering, we modelled the relationship between each cone opsin λ_max_ and a parsimonious microspectrophotometric oil droplet λ_cut_ for Galliformes (Stavenga & Wilts, 2014). The mid-wavelengths of absorbance, *i.e.* where the droplet transmits 50% (or λ_mid_) were inferred for the C, Y and R oil droplet types as follows: For the C-type droplet expressed in SWS2, the established empirical relationship is approximated as C_mid_ = 0.82 x λ_max_SWS1_ + 75, providing C-type λ_mid_ at 430-434 nm (Hart & Vorobyev, 2005); For Y- and R-type droplets expressed in Rh2 and LWS cones, Y_mid_ 515 nm and R_mid_ 560 nm were applied (Hart, 2001, Bowmaker et al., 1997, Hart & Hunt, 2007). Droplet transmission slopes were modelled with 0.2 log unit/10nm, consistent with Hart (2001), with moderate slopes of k = 0.15 for SWS2 and k = 0.12 for Rh2 and LWS. The plotted normalized effective photoreceptor sensitivity range is the product of opsin absorbance x droplet transmission, attenuated relative to opsin absorbance (Dataset 3).

The exponentially decaying visual pigment absorbance spectrum allows to calculate the upper limit of cone sensitivity, sensitivity which is not extended by lateral filtering alone (Bernard, 1979) and can be obtained from the visual Govardovskii template (Govardovskii et al., 2000b) at the wavelength intercepting 5% sensitivity.

### (g) Reflectance spectrum of visual marker devices

Reflectance measurements were carried out using a high-performance spectrophotometer, Cary 5000 Ultraviolet-Visible-Near Infrared (UV-VIS-NIR) spectrophotometer (Agilent) capturing wavelengths every 1 nm from 300 to 780 nm using a photomultiplier tube detector. A reference blank (spectralon) was used for baseline before each measurement. Detection markers installed in ski resort stations were sampled and their characteristics are described in Table S1. Most materials were opaque, therefore not transmitting light rays and preventing measures of light transmission. Two types of reflectance were measured: i) *diffuse reflectance*, in which the incident light is scattered in many directions, and ii) *combined diffuse and specular reflectance*, which includes both scattered and reflected components. Specular reflection corresponds to the direct, angle-dependent reflection and is generally more perceptible at greater distances than diffuse reflection. We measured each component separately, yet in most optical situations, both diffuse and specular components contribute simultaneously to the reflected signal.

To quantify the achromatic contrast between different materials of the same object, we computed Michelson contrast values for the mean spectral reflectance of each material surface. For each sampled area, we averaged the reflectance spectrum over the full measurement range (300–780 nm), yielding a single value representing overall reflectance intensity. The Michelson contrast between two surfaces R_1_ and R_2_ was calculated as:

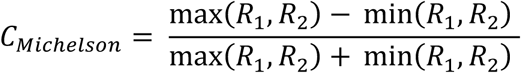

This metric provides a relative measure of contrast that is independent of absolute luminance. We then estimated the inverse contrast value (1/C) to compare with the maximum spectral sensitivity of the black grouse. (Datafile S4). Transparent floats are not included in this analysis as they are composed of a single type of material.

## Results

### (a) Black grouse visual fields

The maximum binocular overlap of the male black grouse covered 40° and is positioned 30° above the bill-tip direction (Figure 2A). At rest, the black grouse has a field of view covering 353°, including a frontal binocular overlap of 28°, lateral fields of 162.5° each, and a blind sector of only 7° behind the head (Figure 2B). Therefore, when black grouses fly at the same altitude as an object (*e.g.* a visual marker), the minimal spacing between two adjacent markers for them to be visible to the black grouse is d = 2*32 *tan(28°) ∼16 m. The vertical extent of the binocular field in the median sagittal plane covers 170° (Figure 2C), extending from 0° to 100° in between the head tip (0°) and the bill-tip (100°), 40° behind the head tip, and 30° below the bill-tip plane (100° -140°) (Figure 2C). At the head tip (0°), the binocular field has a width of 6°.

**Figure 2.**
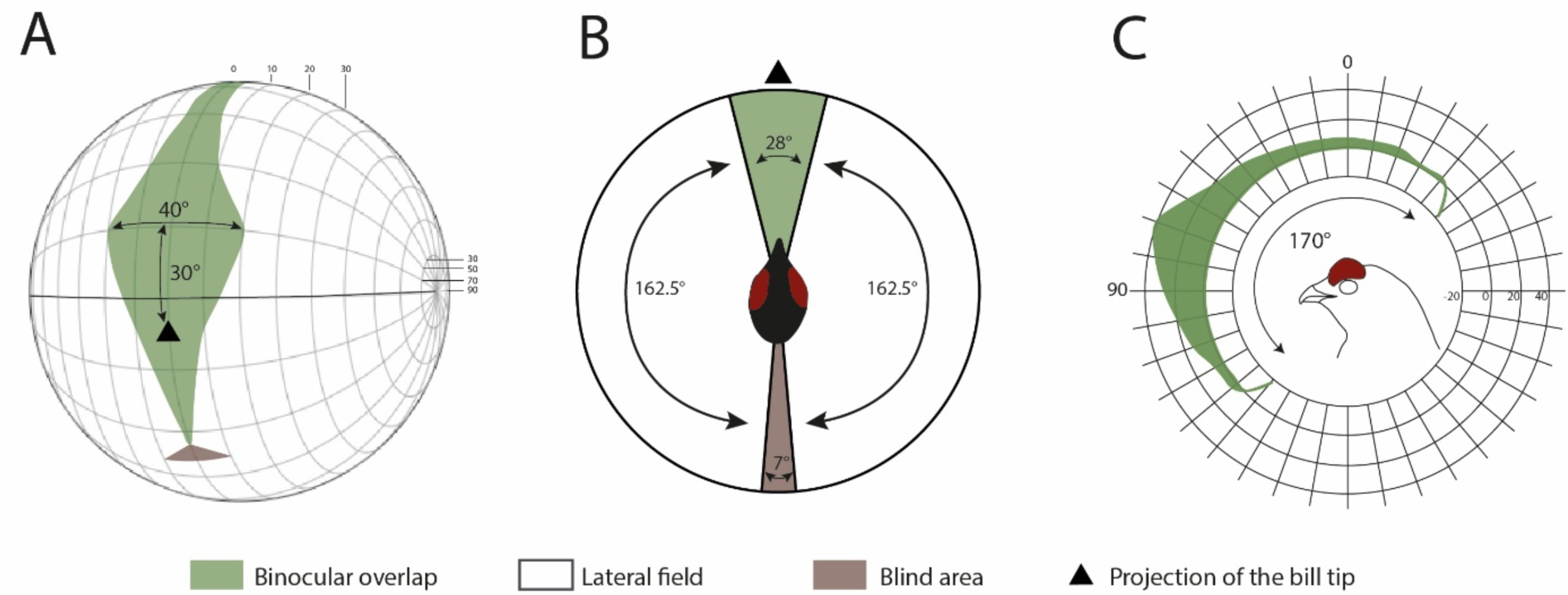
Visual fields of the black grouse. (**A**) Orthographic projection of the retinal field boundaries from both eyes. A latitude and longitude coordinate system is used with the equator aligned vertically in the median sagittal plane. The bird head is positioned at the centre of a schematic globe, and the projection of the bill tip corresponds to the black triangle. The grid lines are drawn at 10° intervals in latitude, and at 20° intervals in longitude. (**B**) Section through the horizontal plane corresponding to the bold line on panel A, and showing the binocular overlap (green), lateral fields (white) and blind spot (brown) in black grouse. (**C**) Binocular field as a function of vertical elevation in the median sagittal plane. The bird head orientation is shown with the direction of the bill tip. All values are in degrees, and the grid is at 10° intervals. Values represent the mean visual field for two measured individuals.

### (b) Eye size, visual acuity and detection distance

The measured Corneal Diameter (CD) from close-up photographs was 8.00 mm for both individuals. The estimated Axial Length (AL) was 13.28 mm, which corresponds to a spatial resolution (SR) of 9.81 c/deg. With a SR = 9.81c/deg, the minimal width (*w*) of an object to be detected at 32m is 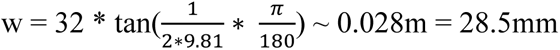.

### (c) Contrast sensitivity

To assess the bird’s ability to distinguish features in its visual environment, we projected stripe patterns with varying contrast and spatial frequencies. Figure 3B shows that contrast sensitivity decreased progressively with increasing spatial frequency—corresponding to a smaller distance to object or increase in object size. Contrast sensitivity also decreased with decreasing spatial frequency—corresponding to a longer distance to object or reduction in object size. This typical contrast sensitivity pattern results in an inverse U-shaped curve (Figure 3B). The estimated optical cut-off frequency, *i.e.* the point at which the stripe pattern becomes too fine for the eye to resolve, was 0.85 cycles per degree (cpd) for individual A and 1.06 cpd for individual B. The average optical cutoff frequency was 0.95 cpd, corresponding to a maximum contrast sensitivity of 16.67 (Figure 3B). This sensitivity corresponds to a Michelson contrast (C_m_) of 6% (*i.e.* contrast sensitivity = 16.67 = 1/C_m_; C_m_ = 1/16.67 ∼ 0.0599 or 5.99%), which means that black grouse can distinguish a signal until it is only 6% brighter than the surrounding background. We then estimated the maximal spatial resolution in a moving environment, that is the detection limit of the visual system when contrast sensitivity is set to 1 (or C_m_ = 1/1 = 1 = 100%), *i.e.* only a black and white grating with 100% contrast would be just detectable. Based on Figure 3B, at a contrast sensitivity of 1, the spatial frequency reaches 1.6 cpd (Figure 3B).

**Figure 3.**
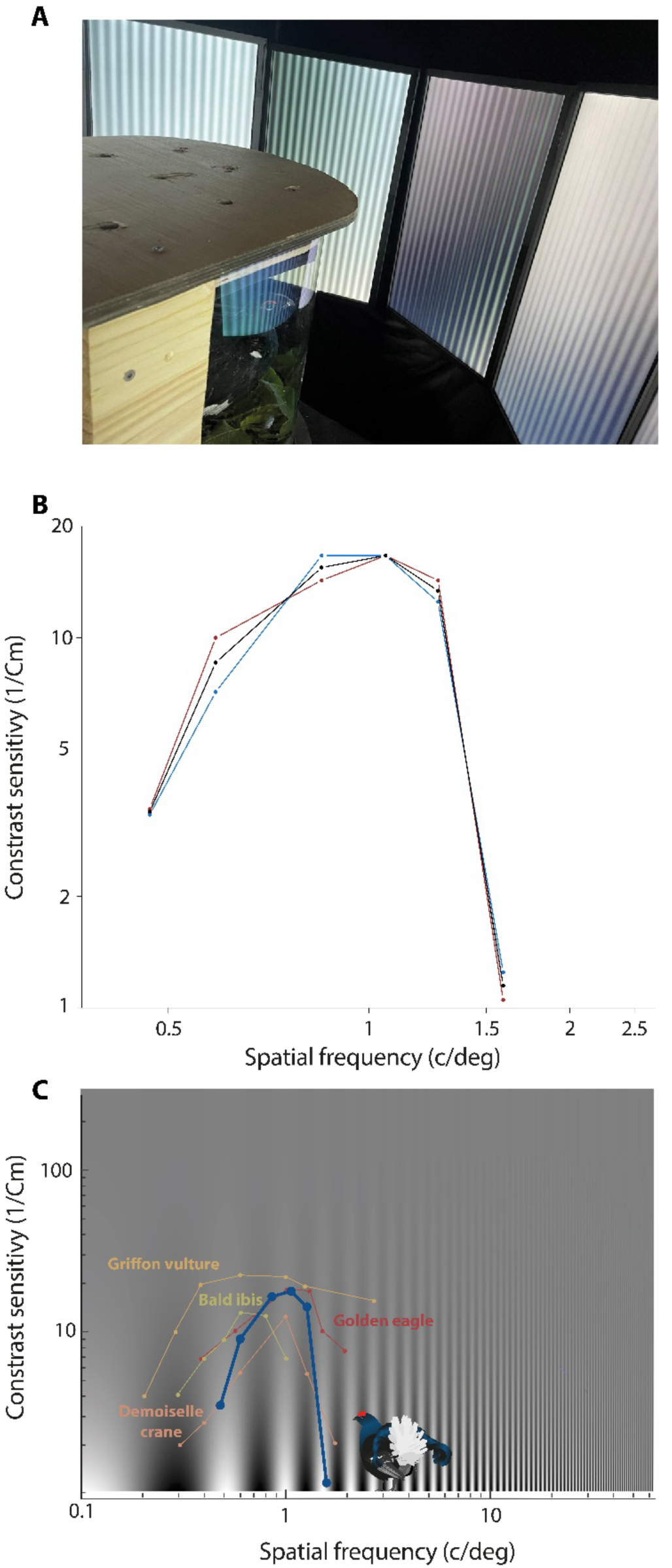
Contrast sensitivity of the black grouse. Contrast sensitivity is defined as the inverse of contrast threshold, as a function of spatial frequency. (**A**) Picture of a black grouse in the apparatus where eyes face monitors displaying gradient stripes. (**B**) Dashed red and blue lines represent contrast sensitivity, expressed as the inverse of Michelson Contrast (1/C_m_) values, for individual A and B. The black curve represents the mean. (**C**) Comparison of the contrast sensitivity between the black grouse (blue curve) and representative species also prone to collisions based on values from Blary et al. (2024). Coloured curves correspond to the following species: red, Golden eagle; yellow, Griffon vulture; green, Bald ibis; orange, Demoiselle crane. Scales in B and C are logarithmic.

### (d) Opsin absorbance

We determined the range of absorbance of the four cone opsins (SWS1, SWS2, Rh2, LW) expressed in the black grouse retina as determined via RNA expression profiling. Active rhodopsin complexes were reconstituted *in vitro* by addition of 11-*cis*-retinal, purified and analysed by ultraviolet-visible spectroscopy to determine their absorbance profiles. These analyses showed that the black grouse SWS1 and SWS2 opsins absorb maximally at λ_max_ of 393 ± 2nm (Figure 4A), and λ_max_ of 436 ± 3nm (Figure 4B), respectively. We then measured the absorbance of black grouse Rh2 and LW, which displayed maximum sensitivity to light at 482 nm ± 2.5 nm (Figure 4C) and 545 ± 2.5 nm (Figure 4D), respectively. Black grouse cone opsins thus perceive wavelengths from ca. 350 nm to 643 nm (Figure 5A) when applying a visual template fit with a 5% sensitivity cut-off for the lower and upper absorbance shoulders of SWS1 and LW, respectively.

**Figure 4:**
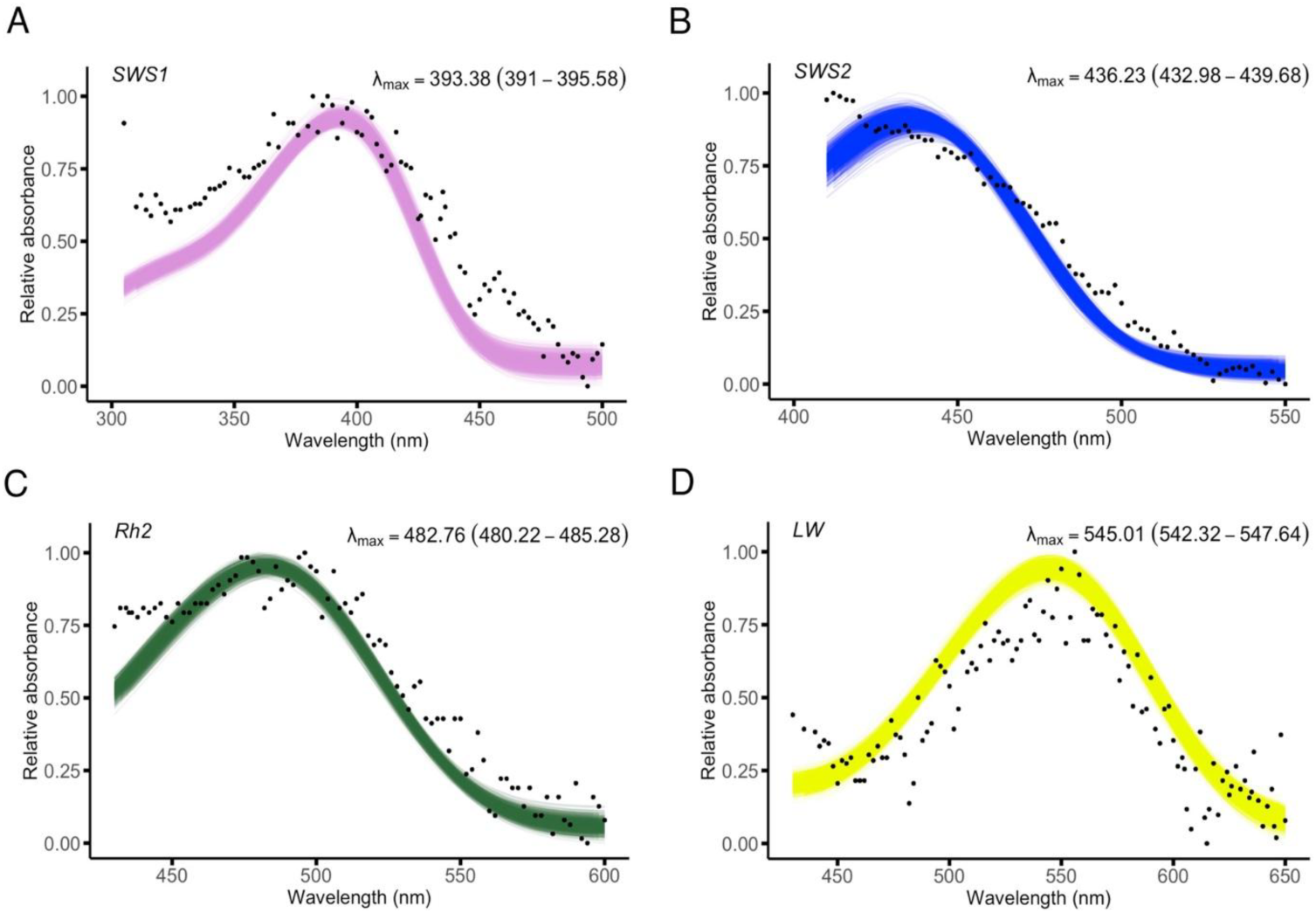
Absorbance range of *Lyrurus tetrix* visual cone opsins purified in complex with 11,*cis*-retinal. (A-D) The black dots represent the average absorbance across replicates at every wavelength, whereas the lines represent the curve-fitting of individual bootstrap replicates. (A) SWS1 (n = 7), (B) SWS2 (n = 6), (C) Rh2 (n = 3), (D) LW (n = 7), where n is the number of measurements of protein aliquots with active rhodopsin complexes. For each rhodopsin, relative absorbance data are fitted to a visual template with polynomial function analyses computed in R to obtain the estimates of lambda max following 1,000 bootstrap replicates. Values in parentheses represent the lower and upper bounds of the 95% confidence intervals. Accompanying data are provided in Dataset S2.

**Figure 5:**
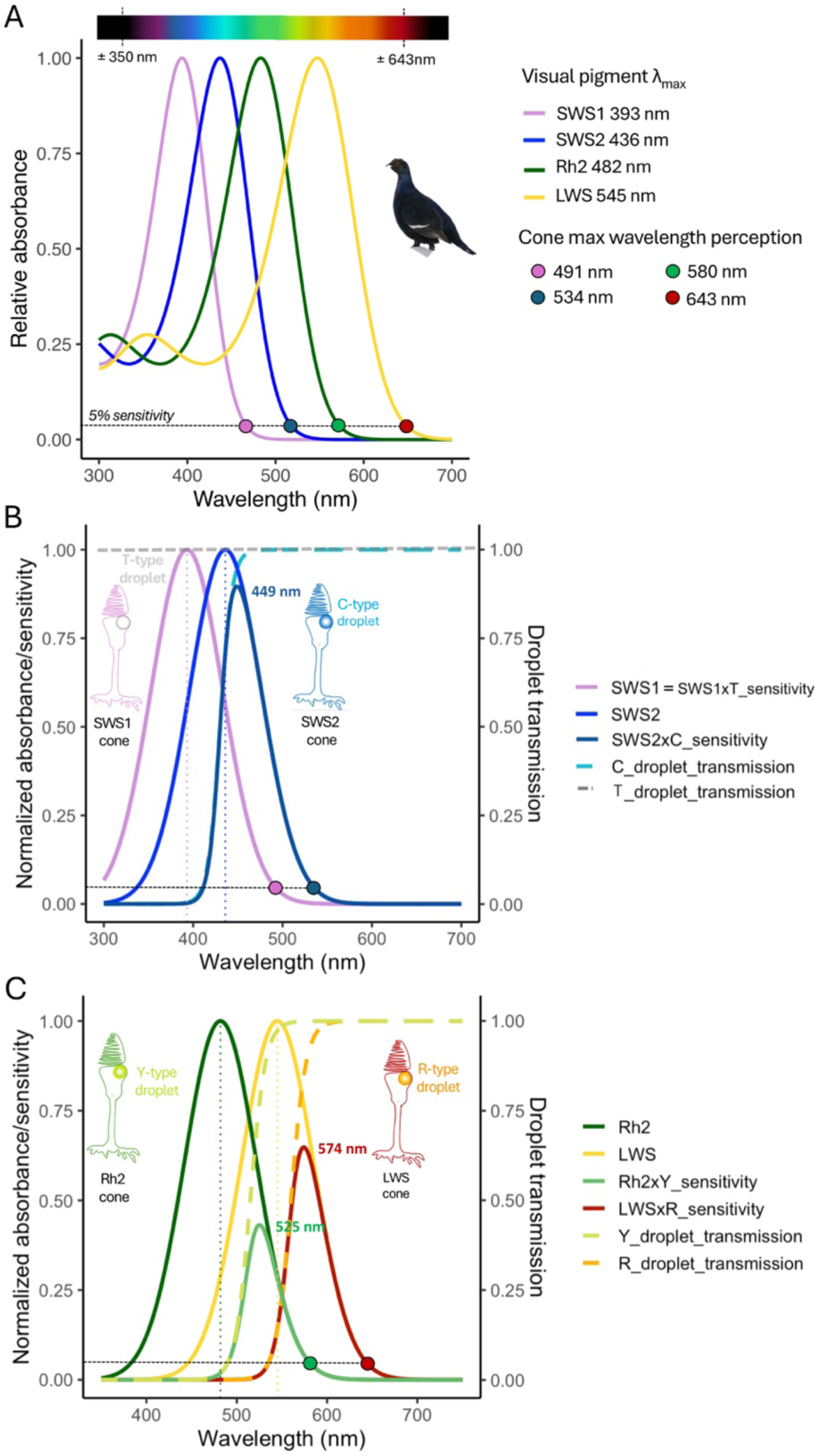
Visual opsin pigment absorbance range and modelled spectral sensitivity in the black grouse. (**A**) The visual pigment absorbance range confers wavelength perception from ca. 350 nm to 643 nm, where opsin absorbance lowers to 5% sensitivity (dashed horizontal grey line). (**B**) Theoretical effective SWS cone spectral sensitivity is calculated as SWS * oil droplet sensitivity. SWS1 cones have transparent T-type oil droplets that transmit light (T = 1) and the effective SWS1 cone spectral sensitivity is similar to the opsin-based absorbance range. In SWS2 cones, C-type droplets filter short wavelengths under 430 nm, following the relationship established by Hart and Vorobyev (2005), modelled as a likely shift of the SWS2 peak sensitivity to approximately 449 nm. (**C**) Theoretical effective Rh2 and LW cone spectral sensitivities. Y- and R-type droplet transmission λ_mid_ values vary moderately across avian species (Y-type 505-515 nm, R-type: 585-613 nm) and estimates of Y-λ_mid_ = 515 nm and R-λ_mid_ = 560 nm are used here, based on known microspectrophotometry average values in other Galliformes such as *Gallus gallus* (Table S4) (Hart, 2001), parsimoniously placing the range of effective cone sensitivity from 525 to 580 nm for Rh2 and from 574 to 643 nm for LWS. Modelling scripts are presented as accompanying Dataset S3.

### (e) Modelled spectral sensitivity

Using the empirical visual pigment λ_max_, which provide the UV-visible spectral range and maximal wavelength perception (Figure 5A), we then modelled the effect of oil droplets on theoretical effective photoreceptor sensitivity in SWS, Rh2 and LW cones (Figure 5B-C). In SWS1 cones, spectral sensitivity corresponds to the visual pigment sensitivity and in SWS2 cones, sensitivity is maximized above 449 nm (Figure 5B). In Rh2 and LW cones, Y- and R-type oil droplets tends to shift the resulting maximal cone spectral sensitivity to longer wavelengths, which modelling indicate would likely be maximised above 525 nm and 560 nm, respectively (Figure 5C). Because oil droplet filters do not alter the long-wavelength limb of absorption by the expressed opsin, the maximal wavelength perceived by the LW photoreceptor remains at ∼643 nm, corresponding to the 5% sensitivity cut-off by the descending absorbance shoulder (Figure 5).

Avian SWS1 opsins are divided into ultraviolet (λ_max_ < 385 nm) and violet-sensitive (λ_max_ >385 nm) types (Hauser et al., 2014), and the black grouse visual pigment, based on its λ_max_ is classified in the violet range but nevertheless absorbs a small fraction of UV light (Figure 3, Figure 4A). We aligned and compared SWS1 opsin sequences between the black grouse and additional representative avian species with known SWS1 opsin absorbance, highlighting amino acid residues at positions previously found involved in spectral tuning (Figure 6) (Hauser et al., 2014, Hagen et al., 2023). The black grouse SWS1 opsin lacks the cysteine residue present in all true ultraviolet sensitive opsins with λ_max_ < 370 nm (Figure 6) and share different degrees of sequence similarities at those spectral positions, with the Chicken (λ_max_ 419 nm) and Pigeon (λ_max_ 393 nm) SWS1 opsins.

**Figure 6.**
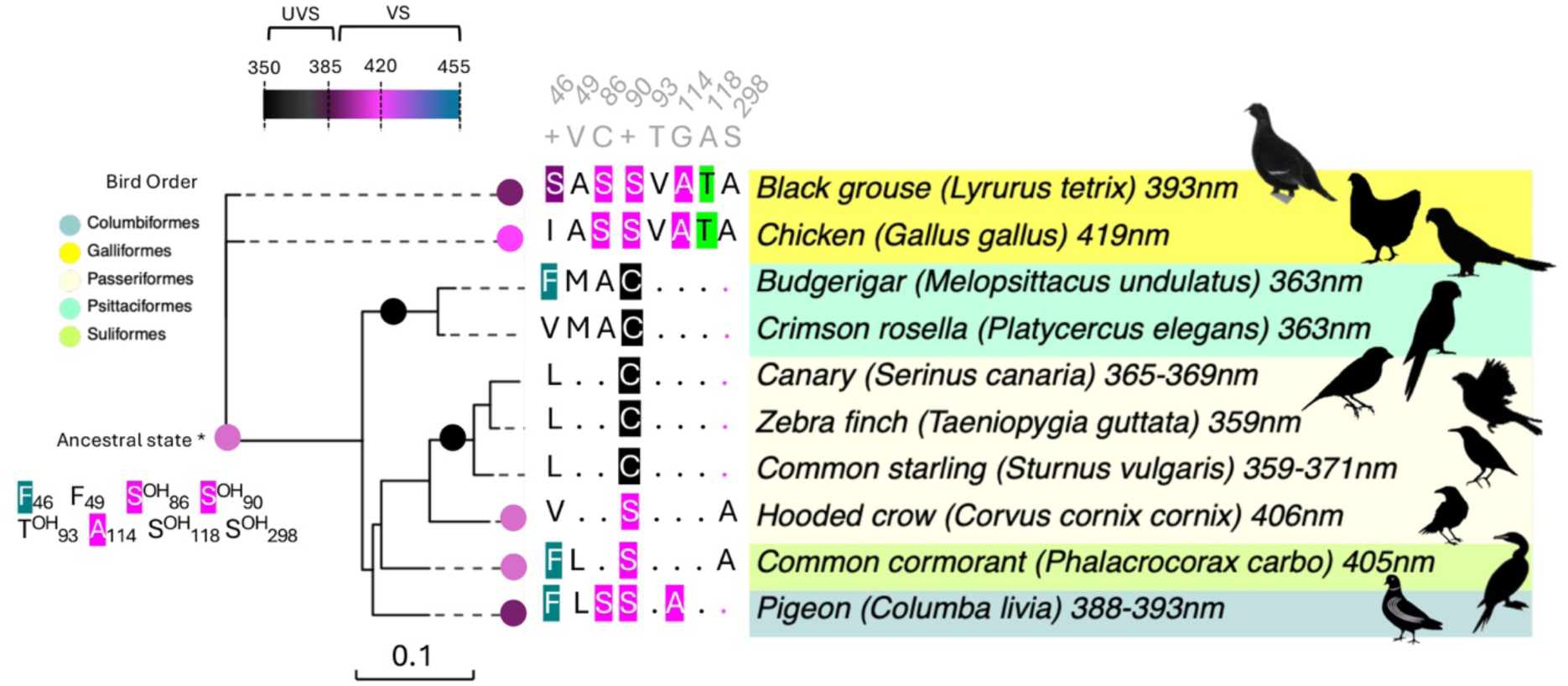
SWS1 opsin phylogeny of representative bird species with known SWS1 visual pigment maximal absorbance (λ_max_). Vertebrate SWS1 pigments are divided into two subtypes (UVS λ_max_ 355-385 nm, VS λ_max_ 385-455 nm) (Hauser et al., 2014) and different circle colours are used to represent UV-shifted (black) to more violet-shifted light. Maximal absorbance peak values measured from purified visual pigments are indicated after each species name and are taken from Hart (2001) and Hart and Hunt (2007). Key amino acid sites and residue substitutions contributing to avian SWS1 maximal sensitivity (reviewed in Hauser et al. (2014)) are indicated along phylogenetic branches, with numbering relative to bovine rhodopsin residue positions (Acc. number AAA30674). The ancestral avian violet SWS1 sensitivity was inferred in Hagen et al. (2023). OH refers to amino acids bearing a hydroxyl group. Dots in the alignment correspond to a conserved residue with the consensus sequence in grey (+ indicates a variable site). Sequence accession numbers: Chicken NP_990769, Budgeriar NP_001298010, Canary NP_001289029, Zebrafinch NP_001070172, Common starling XP_014746280, hooded crow XP_039427152, Common cormorant ABS86975.1, Pigeon NP_001269750.

### (f) Reflectance spectra of visual markers

Visual markers show spectral reflectance within the visible spectrum, and a few materials show reflectance in the ultraviolet range below 400 nm and long-wavelength light above 750 nm (Figure 7, Figures S1 to S8). For most materials except the polished stainless-steel plate (Figure. S7), the specular and diffuse reflection components are nearly identical (Figures S1 to S8). Due to the variety of existing designs, some materials show relative reflectance values exceeding 100% owing to fluorescent elements.

**Figure 7.**
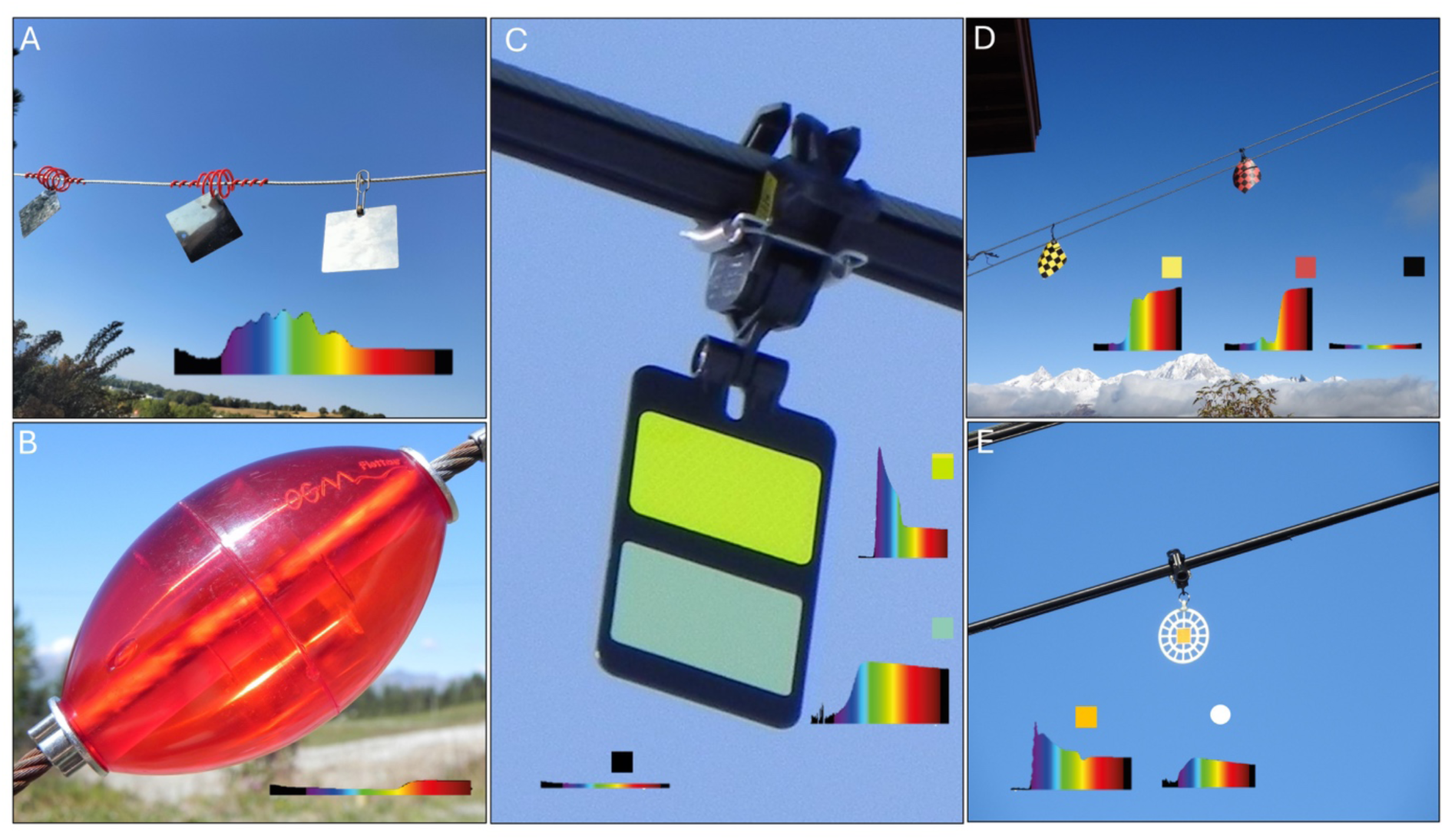
Examples of aerial cables equipped with visual markers and their diffuse reflectance profiles. (see also Table S1 for more designs, and Figures S1-S8 for a comparison between diffuse and specular reflectance). Photographs of (**A)** a *stainless-steel plate* (8 x 8 cm) installed on aerial cables transporting explosives in the context of avalanche control. (**B)** a *transparent float* (7.5 x 4.5 cm) installed on the safety rope of ski lifts. (**C)** a *black firefly* (15 x 9 cm) installed on chairlifts. (**D)** a patterned *flag* (16 x 16 cm) installed on aerial cables for transporting explosives in the context of avalanche control. (**E)** a white orange *Birdmark* (diameter 13.5 cm) installed on chairlifts and safety ropes of ski lifts. Reflectance spectra are measured from 300 nm to 780 nm. Reflectance within the visible range for a human eye is coloured from violet (400 nm) to red (750 nm). Reflectance outside the visible range for a human eye (> 400 nm and < 750 nm) are coloured in black.

### (g) internal contrasts of visual markers

For most visual markers, internal achromatic contrast between different materials was generally below the black grouse maximal sensitivity contrast of 16.67 (Figure 8), in other words most of these visual features have enough brightness contrast to be perceived by the black grouse visual system. Reversely, specific combinations of adjacent materials may not meet the necessary contrast threshold for detection by black grouse. The orange retroreflectors on *Birdmarks*, but not the yellow one, exhibited a low achromatic contrast (*i.e.* a high 1/Michelson contrast) with the paddle, contrast which falls outside of the limits of sensitivity for the black grouse. Similarly, the achromatic contrast observed for the yellow retroreflector and the paddle on the *White Crocfast* appears outside the contrast detection limits. For the *White Firefly*, the white beacon lacked sufficient contrast with the adjacent yellow retroreflector. For the *Black Firefly*, both the yellow and phosphorescent retroreflectors showed low achromatic contrast; however, this is partially mitigated by the presence of the black beacon, which provides a degree of visual separation between the elements.

**Figure 8.**
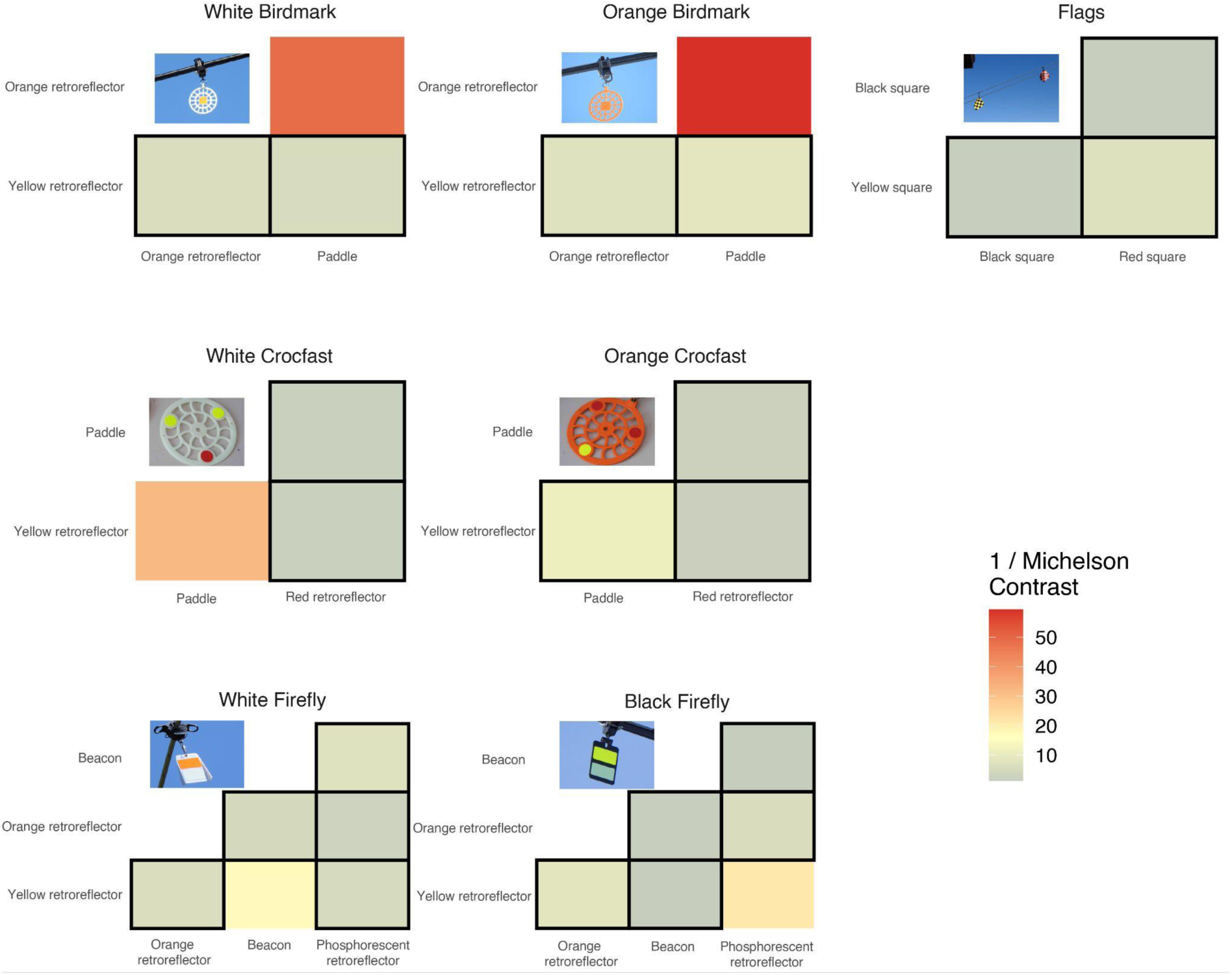
Heatmap representation of internal contrasts in visual markers composed of multiple materials. Contrasts are calculated as the inverse of the Michelson contrast threshold. Squares outlined in black indicate cases where contrasts fall below the maximum contrast sensitivity of the black grouse (1/Michelson contrast <= 16,67), suggesting that the visual system can perceive the difference in brightness between components of the visual marker, which exceeds the smallest brightness difference detectable by the black grouse visual system at its peak Critical Flicker Fusion (CFF) frequency, that is when the visual system is most sensitive to changes in brightness. See also Dataset S4.

## Discussion

Constraints in bird visual perception have been suggested as one of the factors contributing to variable risks of collisions with man-made structures (Martin, 2011a, Martin, 2022, Martin & Banks, 2023). In this study, we examined visual characteristics of the black grouse, specifically its visual fields, contrast sensitivity, corneal size (and the estimated visual acuity), as well as visual pigment spectral range, to gain a better understanding of its multifaceted vision, and discuss how these factors may affect the perception of obstacles and visual markers.

### Visual fields

Avian visual fields reflect ecological differences, such as in navigation and foraging, parental care, or predator detection, and vary significantly among avian species (Martin, 2017b). The visual field configuration of the black grouse shows that this species has an exceptionally extended visual field in both the horizontal and vertical planes. From more than 250 avian species studied to date (Martin, 2017a, Potier et al., 2018a, Cantlay et al., 2023, Potier et al., 2023), only four have a blind visual sector behind the head smaller than that of the black grouse (7°), *i.e.* the Eurasian woodcock *Scolopax rusticola* (Martin, 1994) and three duck species, the Mallard *Anas platyrhynchos*, the Northern pintail *Anas acuta* and the Pink-eared duck *Malacorhynchus membranaceus* (Martin, 1986, Guillemain & Fritz, 2002, Martin, 2007).

In the context of avian collision, both the binocular overlap and vertical extent of the binocular field are important parameters. The binocular overlap is thought to function primarily in the detection of symmetrically expanding optic flow-fields that provide almost instantaneous information on the direction of travel and time-to-contact a target, whereas the vertical extent defines whether birds can see in the direction of their flight path (Martin, 2009, Martin & Portugal, 2011, Martin et al., 2012, Martin, 2017a, Martin, 2022).

The black grouse frontal binocular field covers 28° on the horizontal plane, in agreement with typical avian values (20°-40°) (Martin, 2017b), and has a wide maximal binocular overlap of 40°, located 30° above the eye bill-tip direction (Figure 2A,C), which is similar to other visual ground forager species such as the Southern caracara *Caracara plancus* (Potier et al., 2017). Together, the binocular field of the black grouse is well adapted to its ground foraging behaviour.

The vertical extent of the binocular field allows black grouse to gain a comprehensive visual coverage of the celestial hemisphere, and the ability to see the way ahead irrespective of their head position. This configuration should therefore enable black grouse to detect aerial objects. In Galliformes, however, suborbital ridges develop in adult males (Lindén & Väisänen, 1986) or during the breeding season (Lamond et al., 2025), suggesting that the vertical binocular field may be somewhat reduced in older males, which merit further quantification across development, including comparison with females.

In contrast to the black grouse, many large raptors, such as some eagles, vultures or hawks (Accipitriformes) may fail to see the way ahead (Martin et al., 2012, Portugal et al., 2017, Potier et al., 2018a). Hence, while in flight visualizing the ground below for food, their limited field of view would comparatively provide none or only intermittent information about the flight path ahead (Martin et al., 2012).

Thus the finding that black grouse can detect objects in a wide horizontal and also in the vertical plane is informative to directly optimize aerial man-made structures, by modifying *e.g.* an object’s size or spacing, its internal contrasts or contrast relative to the environmental background, or its spectral properties (colours), and/or using visual markers (Martin, 2022) to improve the detection of object.

### Spatial resolution, visual acuity and detection distance

Avian spatial resolution is highly variable, from as low as 4 cpd in the barn owl *Tyto alba* to as high as 142 cpd in the Wedge-tailed eagle *Aquila audax* (reviewed in Martin (2017a)). The black grouse has a spatial resolving power of 9.81 cpd, which is relatively low. Its visual acuity, estimated from eye size, is nevertheless very similar to that of closely related species in the Order Galliformes, ranging from 9.7 cpd in the Japanese quail *Coturnix japonica* to 13 cpd in the Sharp-tailed grouse *Tympanuchus phasianellus* (Lisney et al., 2012a). Although additional precision may be gained through approaches based on photoreceptor density or behavioural assays, these results show that eye size remains a reliable proxy for spatial resolution (Kiltie, 2000, Caves et al., 2024) and can provide estimates comparable to those obtained with operant conditioning (Potier et al., 2016).

Spatial resolution informs at which distance black grouse can detect objects in optimal environmental conditions, that is i) with no air distortion and a clear sky, ii) for a black object on a white background (*i.e.* high contrast scenario), iii) high daylight level, and iv) with the bird visually fixing the object with its acute centre of vision (Land & Nilsson, 2012, Martin, 2017b, Martin, 2022). Under optimal conditions, the shortest distance at which the black grouse can detect two adjacent objects while in flight is 32 m, a conservative minimal value that may increase depending notably on bird motion, atmospheric turbulence or directional lightning in mountain, conditions all known to affect flight in real time. Considering the visual acuity as a proxy of corneal diameter, trigonometric calculations indicate that, to be detected at 32 m, an object should be of a minimum width of 28.5 mm. This measure is 4 to 5 times larger than the smallest object detectable by an average human eye with its spatial resolution of 40-50 cpd (Martin, 2017b). In addition, in a natural scene, optimal conditions are rarely reached; spatial resolution declines up to 5-fold with decreasing light levels *e.g.* on a cloudy day, or at dusk and dawn (Land & Nilsson, 2012, Martin, 2022). To compensate for this lower resolution, the object width should linearly be increased 5 times, therefore reaching a minimum width of 142.5mm, a width currently only met by *flag* designs (15 cm x 15 cm, Figure 7D, Table S3).

### Contrast sensitivity

The maximum contrast sensitivity of the black grouse (16.67) is in the middle range compared to various birds, *e.g.* from 4.8 in the Bourke’s parrot *Neopsephotus bourkii* to 31 in the American kestrel *Falco sparverius* (Hirsch, 1982, Lind et al., 2012, Blary et al., 2024). Values in birds are typically lower than contrast sensitivity observed in other vertebrates such as most rodents (30) or human (close to 200) (De Valois et al., 1974, O’Carroll & Wiederman, 2014). Since the black grouse maximum contrast sensitivity is ∼10x lower compared to a human eye, a dark aerial cable or a dark lift structure against a dark green forest background, or a visual marker against a snowy background (Figure 1) would be more difficult for a black grouse eye to detect compared to a human eye. Consequently, choosing markers with high internal object contrasts (Figure 8), as well as high contrast with the surrounding environment (Martin, 2022) should further aid black grouse in detecting aerial infrastructures under variable light conditions.

### Spectral range

The spectral characteristics of the black grouse visual opsin pigments align with avian pigment sensitivities (reviewed in Hagen et al. (2023), and together its four avian cone opsins absorb wavelengths of light spanning ultraviolet to red from ca. 350 to 643 nm. Whereas transmission properties of cone oil droplets in the black grouse are not yet known, models based on other galliform species are informative to infer the black grouse effective spectral sensitivity (Figure 5). In SWS1 cones, transparent T-type oil droplets do not absorb light (Hart & Vorobyev, 2005), so maximal spectral sensitivity aligns primarily with the SWS1 visual pigment λ_max_. Differences in absorbance characteristics of the avian ocular media can also tune UV sensitivity since the cornea typically also filters a portion of UV light, albeit with large interspecific variation among species *i.e.* low and high cut-offs vary between ∼315 and 370 nm (reviewed in Hart and Vorobyev (2005)). The violet-sensitive black grouse SWS1 pigment, with an absorbance peak around 393 nm, would likely be capable of detecting some signals in the UV range, even in the case of a relatively high corneal filtering that would absorb short-UV wavelengths. Behavioural research has shown that black grouse preferentially choose UV-reflecting berries (Siitari et al., 2002), suggesting that black grouse SWS1 cones indeed enable perceiving at least some long ultraviolet wavelengths, although further work characterizing the UV absorbance of black grouse lenses is needed. Differences in opsin λ_max_ are largely contributed by molecular variation at tuning spectral sites and have been shown to influence colour perception in vertebrates (Hauser et al., 2014, Hagen et al., 2023). The ecological consequences of small variations within avian violet-sensitive pigments however remain to be studied in more detail.

Together with the black grouse SWS2 pigment absorbance, the modelled spectral sensitivity of SWS2 cones (λ_max_ 449 nm) after C-filtering aligns closely with in vivo measurements in related galliform bird species, such as the Japanese quail (*Coturnix japonica*, λ_max_ ∼ 450 nm; (Bowmaker et al., 1993)) or domestic chicken (*Gallus gallus*, λ_max_ ∼ 450 nm; (Bowmaker et al., 1997)), indicating that black grouse SWS2 cones are primarily sensitive to wavelengths above 400 nm up to 534 nm. Rh2 cones are sensitive from above 500 nm to 580 nm with maximal spectral sensitivity predicted at 525 nm after Y-filtering. This wavelength effectively defines the upper limit for opponency between Rh2 and LWS cones and therefore sets the functional limit for chromatic colour discrimination, which based on our values of pigment absorbance and sensitivity modelling in black grouse is restricted to the blue–yellow range. The LWS cones have a predicted spectral peak sensitivity at 574 nm after R-filtering, and an upper sensitivity limit at 643 nm, defined by the LW pigment, allowing black grouse to perceive the brightness of orange and red objects. However, because the chromatic contrast is limited by Rh2 sensitivity, most wavelengths above ∼600 nm are likely not distinguished as colours, although achromatic (brightness) perception remains effective up to the LWS 5% sensitivity cut-off at ∼643nm. Compared to humans, whose MWS and LWS cones are shifted to 535 nm and 564 nm (Kelber, 2019), respectively, black grouse are thus less capable of chromatically discriminating red–orange hues.

### Detection abilities of visual markers and considerations for future optimisation

Across a large diversity of materials used to build visual markers currently in use in ski areas in the French Alps, we found that all tend to reflect visible wavelengths of light in the range of perception of the black grouse. Wavelengths shorter than 400 nm are rarely reflected overall, and some of the red visual markers include long-wavelength reflectance above 650 nm, presumably beyond the limits of black grouse perception.

For the black grouse, a marker may thus not benefit from chromatic contrasts above 600nm (*i.e.* adjacent yellow and orange, or orange and red patterns would be inefficient). Reversely, visual markers would provide strong chromatic contrasts if they reflect light from 400 nm to 580 nm (*i.e.* from violet/blue to yellow/orange spectra), and potentially also achromatic signals from long-UV reflectance (∼370-400 nm), although the latter would need to be tested experimentally. In the Sandhill Cranes *Grus canadensis*, for which the SWS1 pigment of a closely related *G. americana* species has a λ_max_ at 404 nm (Porter et al., 2014), night collisions with power lines decreased by 98% after introducing UV lamps emitting short wavelengths (380-390 nm) on power poles (Dwyer et al., 2019), suggesting some UV sensitivity, similarly to the black grouse. Although black grouse are mainly diurnal and their flight behaviour differs from cranes, increased UV-contrast could be considered in future modelling or field studies to assess their perception against natural backgrounds (Baasch et al., 2022), and under varying natural environmental light.

Chromatic contrasts and spectral sensitivity are thus important to consider in collision contexts (Fernández-Juricic et al., 2011, Zheng et al., 2022), and greater differences in the reflectance spectra of spatially adjacent materials correspond to higher internal contrasts, which several visual markers readily possess, such as the *crocfast*, *flags* and *firefly* designs (Figure 8). In line with our observations, high-contrast striped coloured patterns have recently been shown to reduce bird approaches on wind turbines, interestingly possibly due to a resemblance with unattractive aposematic patterns (Hancock et al., 2025).

In addition to chromatic contrasts, luminance is a highly reliable parameter across different light levels (Land & Nilsson, 2012) and at longer distance because the achromatic spatial resolution is generally higher than the chromatic one (Lind & Kelber, 2011, Potier et al., 2018b). A high internal contrast would both increase visual discrimination against different types of background and improve detection under variable light conditions. Finally, since birds rely on their binocular field to assess the direction of travel and time to contact with an object in their flight path, two adjacent visual makers should be just visible in the bird’s binocular field (Martin, 2022). In the case of the black grouse, adjacent markers should be placed no further apart than 17 m and have a minimal width of 142 mm to ensure reliable continuous perception while in flight. This study provides valuable sensory insights applied to the issue of black grouse collisions on aerial anthropogenic infrastructures. To robustly link these visual sensory mechanisms to collision-related mortality will ultimately require field-based ecological validation of cable detection behaviour with and without visual markers.

## Ethics

Ethical approval for the handling and restraint of birds was granted under the permit number APAFIS# 31775-31775-2021041311196892 v5.

## Data accessibility

All data are available as electronic supplementary materials (SI file) accompanying this manuscript or can be accessed via Figshare https://figshare.com/s/dc412ea22f2eca8ff869 (Datasets 1 to 4).

## Acknowledgements

The authors thank Frank Grosemans for permission to access black grouse individuals, Pascal Barla for expertise on spectral rays, the Fédération des Chasseurs de Savoie (FDC73), Catherine Desnos and Pascale Malchiodi for logistical and technical assistance, and two anonymous reviewers for their insightful criticisms which helped us improve the study. Computational analyses were run on the GIGA Cluster at the University of Liège.

