## Supplementary material for "Exploring the visual system of the Black Grouse (*Lyrurus tetrix*): Combining experimental and molecular approaches to inform strategies for reducing collisions"

### Supplementary tables

Table S1. Visual marker devices

Table S2. Contrast sensitivity

Table S3. Oligonucleotide primers

Table S4. Modelling of effective cone sensitivity

### Supplementary figures

Figure S1. Diffuse and specular reflection of Birdmarks

Figure S2. Diffuse and specular reflection of Crocfast

Figure S3. Diffuse and specular reflection of Flags

Figure S4. Diffuse and specular reflection of white Fireflies.

Figure S5. Diffuse and specular reflection of black Fireflies.

Figure S6. Diffuse and specular reflection of Floats

Figure S7. Diffuse and specular reflection of stainless-steel plates

Figure S8. Diffuse and specular reflection of spirals and fluorescent tubes

**Table S1.Visual marker devices.**

| Type | Photograph |
|---|---|
| *Spirale avifaune*<br>Installation on the multipair of chairlifts<br>product retired<br>Size: 30 cm x 4 cm | 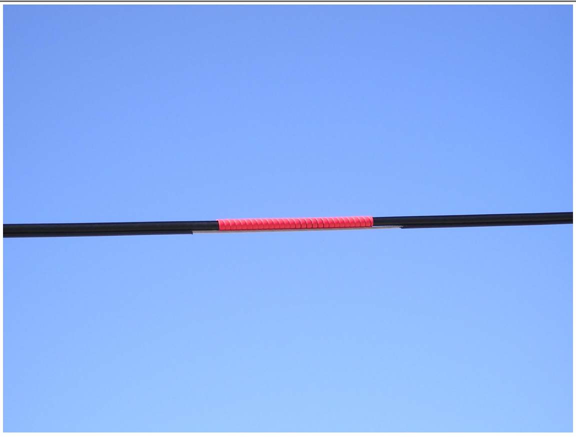 |
| *Transparent float*<br>Installation on safety rope of ski lifts<br>Size: 7.5 cm X 4.5 | 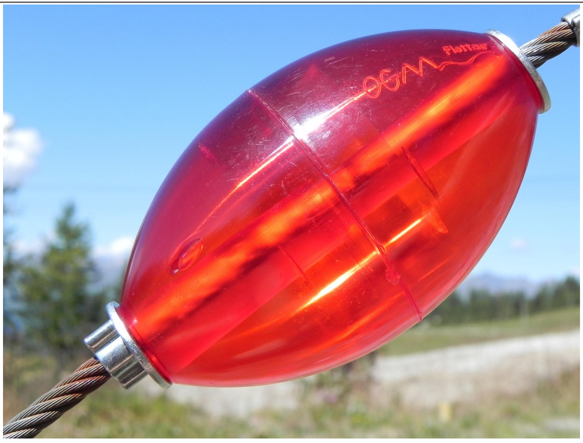 |
| *Opaque float* (OGM Flotteur)<br>Installation on safety rope of ski lifts<br>Size: 7.5 cm X 4.5 | 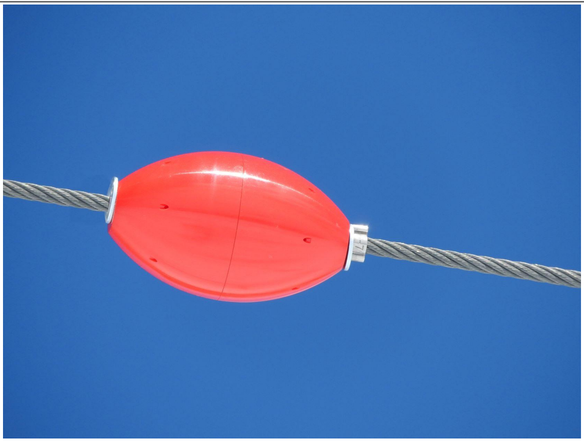 |

| | |
|---|---|
| *Fluorescent tube*<br>(used to equip guy cables)<br>Size: 15 cm x 3.5 cm | 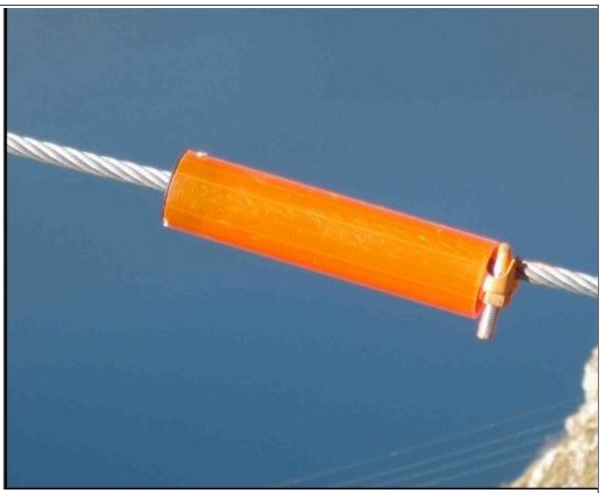 |
| *Shiny stainless-steel plate*<br>To equip aerial cables for transporting explosives<br>Size: 8 x 8 cm | 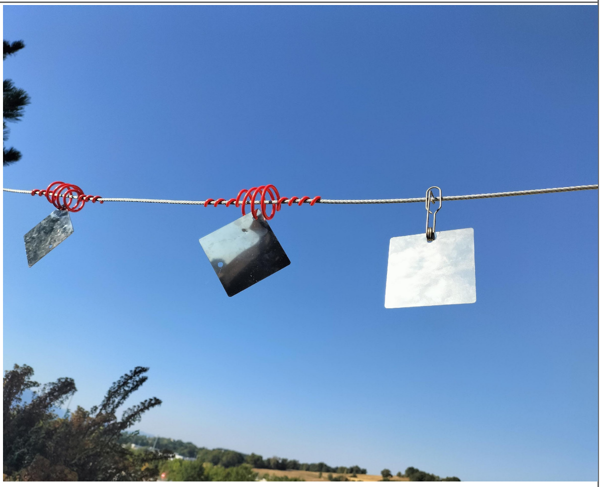 |
| *Matte stainless-steel plate*<br>Size: 8 x 8 cm | 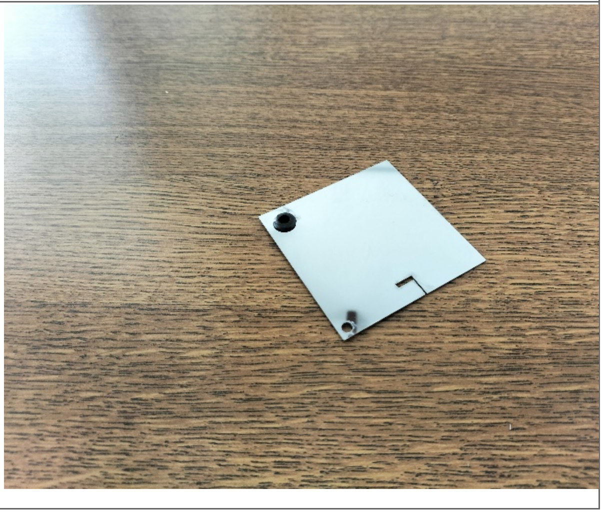 |

| | |
|---|---|
| *White Birdmark*<br>(stores light that is reflected at night)<br>Installation on the multipair of chairlifts, gondola and on safety rope of ski lifts<br>Installation recommendation:<br>Alternation White Birdmark and Orange Birdmark<br>Size: Racket diameter = 13.5 cm | 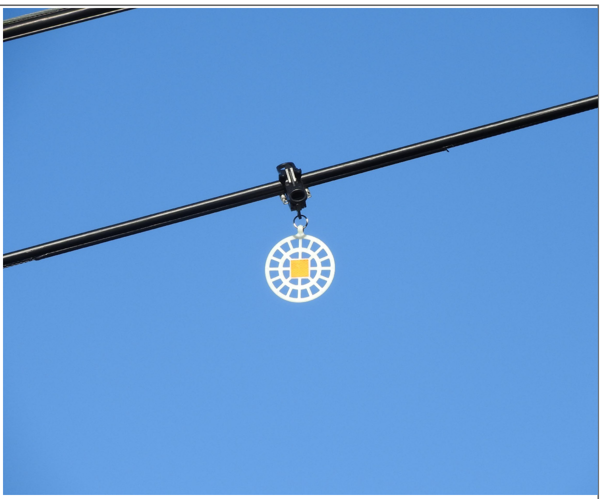 |
| *Orange Birdmark*<br>Installation on the multipair of chairlifts, gondola and on safety rope of ski lifts<br>Installation recommendation:<br>Alternation White Birdmark and Orange Birdmark<br>Size: Racket diameter = 13.5 cm | 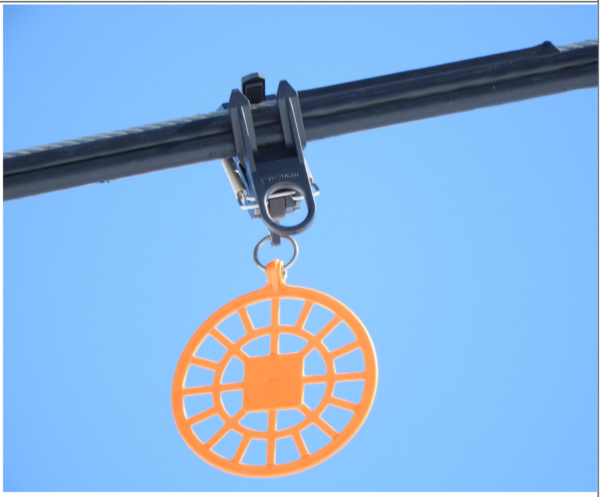 |
| *Crocfast (test)*<br>Size: Racket diameter = 13.5 cm | 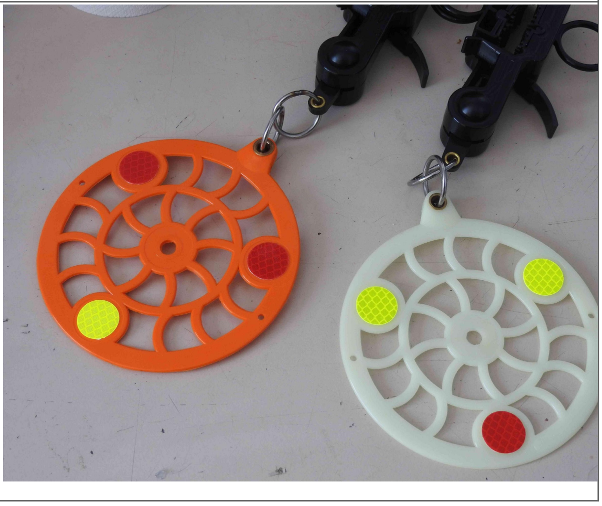 |

| | |
|---|---|
| *Black Firefly*<br>Installation on the multipair of chairlifts and gondola<br>Size: 15 cm x 9 cm | 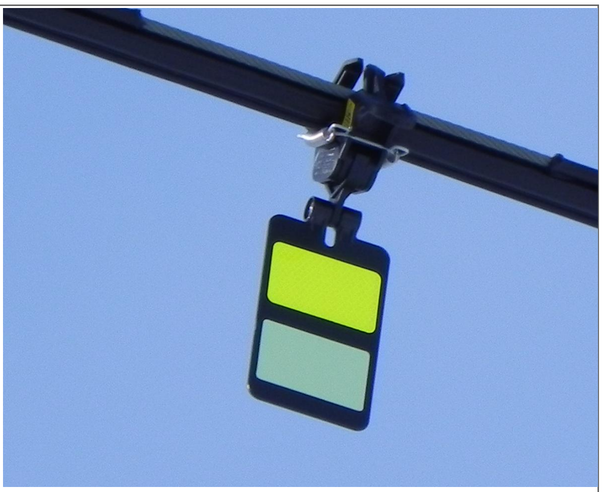 |
| *Firefly*<br>equipment for power lines<br>Size: 15 cm x10.5 cm | 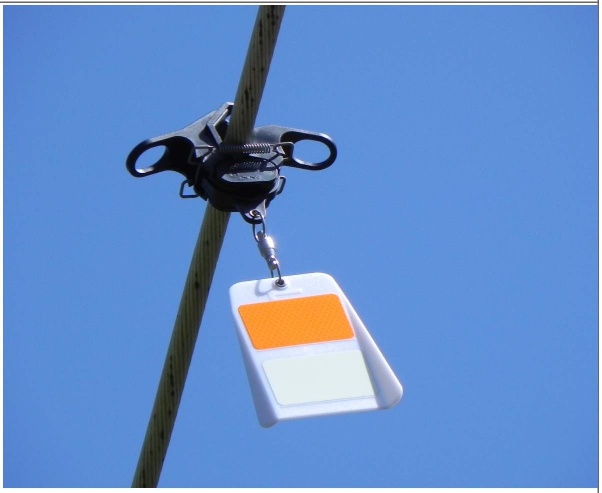 |
| *Flag*<br>To equip aerial cables for transporting explosives<br>Size: 16x16 cm | 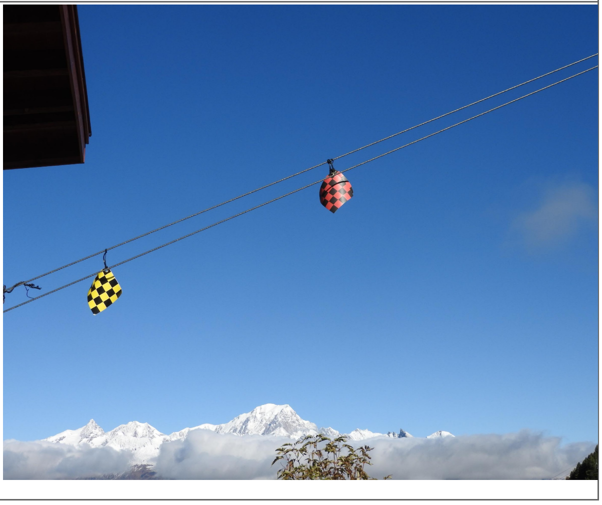 |

**Table S2. Contrast sensitivity.** Steps used to calculate contrast sensitivity, contrast values associated with the vertical sinusoidal achromatic gratings, and the average luminance of the screens.

| Step | Michelson Contrast | Inverse Michelson Contrast | Average luminance ($cd/m^2$) |
|---|---|---|---|
| 127 | 1.00 | 1.00 | 130.26 |
| 120 | 1.00 | 1.00 | 124.79 |
| 110 | 0.99 | 1.01 | 116.42 |
| 100 | 0.97 | 1.03 | 107.09 |
| 90 | 0.94 | 1.07 | 99.86 |
| 80 | 0.90 | 1.11 | 93.51 |
| 70 | 0.85 | 1.18 | 87.80 |
| 60 | 0.78 | 1.29 | 82.73 |
| 50 | 0.69 | 1.45 | 77.88 |
| 45 | 0.64 | 1.57 | 75.36 |
| 40 | 0.58 | 1.71 | 73.18 |
| 35 | 0.52 | 1.92 | 71.24 |
| 30 | 0.46 | 2.17 | 69.79 |
| 29 | 0.45 | 2.24 | 69.33 |
| 28 | 0.43 | 2.31 | 68.97 |
| 27 | 0.42 | 2.39 | 68.53 |
| 26 | 0.40 | 2.47 | 68.28 |
| 25 | 0.39 | 2.55 | 67.95 |
| 24 | 0.38 | 2.65 | 67.89 |
| 23 | 0.37 | 2.74 | 67.31 |
| 22 | 0.35 | 2.88 | 67.21 |
| 21 | 0.33 | 2.99 | 67.06 |
| 20 | 0.32 | 3.13 | 66.67 |
| 19 | 0.30 | 3.29 | 66.57 |
| 18 | 0.29 | 3.45 | 66.34 |
| 17 | 0.27 | 3.64 | 66.09 |
| 16 | 0.26 | 3.84 | 65.88 |
| 15 | 0.24 | 4.10 | 65.57 |
| 14 | 0.23 | 4.39 | 65.33 |
| 13 | 0.21 | 4.71 | 65.21 |
| 12 | 0.20 | 5.07 | 64.98 |
| 11 | 0.18 | 5.50 | 64.71 |
| 10 | 0.17 | 6.04 | 64.62 |
| 9 | 0.15 | 6.62 | 64.49 |
| 8 | 0.13 | 7.50 | 64.13 |
| 7 | 0.12 | 8.51 | 63.96 |
| 6 | 0.10 | 9.62 | 63.78 |
| 5 | 0.09 | 11.46 | 63.57 |
| 4 | 0.07 | 13.89 | 63.50 |

| | | | |
|---|---|---|---|
| 3.5 | 0.06 | 16.12 | 64.05 |
| 3 | 0.05 | 18.32 | 63.54 |
| 2.5 | 0.05 | 21.70 | 64.04 |
| 2 | 0.04 | 26.73 | 63.35 |
| 1.5 | 0.03 | 35.39 | 63.89 |
| 1 | 0.02 | 51.29 | 63.23 |
| 0.5 | 0.01 | 82.54 | 63.76 |

Table S3. Oligonucleotide primers

| ORF primers | Sequence (5' - 3') | ORF size (bp) |
|---|---|---|
| *s,forward; as, reverse* | *nucleotides with underlined capital letters: restriction sites; nucleotides in italics: extra bases for restriction digestion; nucleotides in bold: Kozak sequence* | |
| Lte-SWS1-ORF-HindIII-s | *ggtgcc*AAGCTT**GCCACC**ATG**G**CCTCGGACGACGACTTCTACCT | 1,044 |
| Lte-SWS1-ORF-BsiWI-as | *cggccg*CGTACGAGTGGGGCTGACCTGGCTGG | |
| Lte-SWS2-ORF-AflII-s | *ggtgcc*CTTAAG**GCCACC**ATG**G**ACCCCCCTCGCCCCC | 1,071 |
| Lte-SWS2-ORF-BsiWI-as | *cggccg*CGTACGGGCGGGCGAGACGGAGG | |
| Lte-Rh2-ORF-HindIII-s | *ggtgcc*AAGCTT**GCCACC**ATG**G**GACAGAAGGTATCAATTTTTA | 1,068 |
| Lte-Rh2-ORF-BsiWI-as | *cggccg*CGTACGTGCAGGTGATACTTGGCTGGAGG | |
| Lte-LW-ORF-HindIII-s | *ggtgcc*AAGCTT**GCCACC**ATG**G**AGGAGTGGGAGGCGGCG | 1,089 |
| Lte-LW-ORF-BsiWI-as | *cggccg*CGTACGGGCGGGCGAGACGGAGGA | |
| **pcDNA5-FLAG_T2A-mruby2 construct verification primers** | | |
| pcDNA5-s | GCTGTTTTGACCTCCATAGAAGA | - |
| pcDNA5-as | TAGAAGGCACAGTCGAGG | - |
| pLenti-s | ACCGCATGTTAGCAGACTT | - |
| mRuby-as | CGGCGGCTTAAACCTTATCGTCG | - |

Table S4. Modelling of effective cone sensitivity

| Cone type | Opsin $\lambda_{max}$ (nm) | Droplet type | Droplet cut-off $\lambda_{mid}$ (nm) | Effective cone $\lambda_{max}$ (nm) | Upper cone sensitivity (5% threshold) (nm) |
|---|---|---|---|---|---|
| SWS1 | 393 *(a)* | T-type | none | 393 | 491 |
| SWS2 | 436 *(a)* | C-type | ~430 *(b)* | 449 | 534 |
| Rh2 | 482 *(a)* | Y-type | ~515 *(c)* | 525 | 580 |
| LWS | 545 *(a)* | R-Type | ~560 *(c)* | 574 | 643 |

*(a) this study;*
*(b) based on MSP values from Hart and Vorobyev 2005 (DOI: 10.1007/s00359-004-0595-3);*
*(c) based on MSP values from Hart 2001 (DOI: 10.1016/s1350-9462(01)00009-x)*

### Supplementary Figures

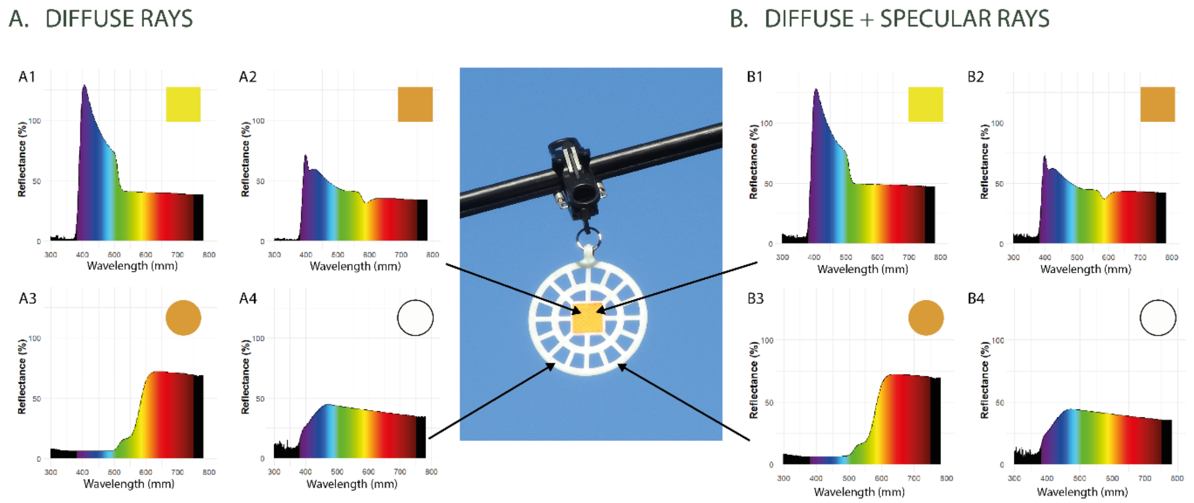


**Figure S1**. **Diffuse and specular reflection of Birdmarks**. A1 and B1 correspond to the yellow retroreflector, A2 and B2 correspond to the orange retroreflector, A3 and B3 correspond to an orange paddle, and A4 and B4 correspond to a white paddle.

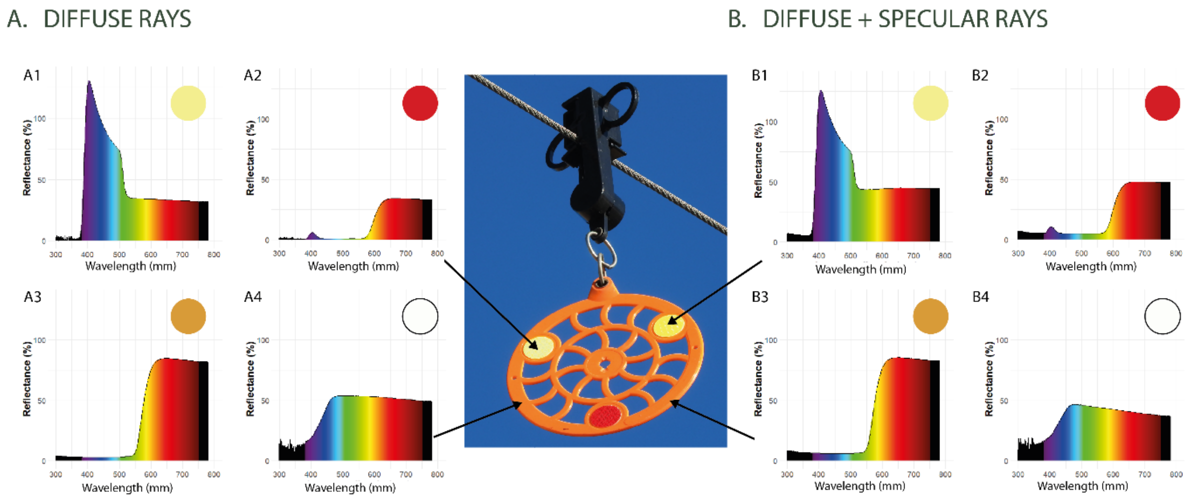


**Figure S2**. **Diffuse and specular reflection of Crocfast**. A1 and B1 correspond to the yellow retroreflector, A2 and B2 correspond to the red retroreflector, A3 and B3 correspond to an orange paddle, and A4 and B4 correspond to a white paddle.

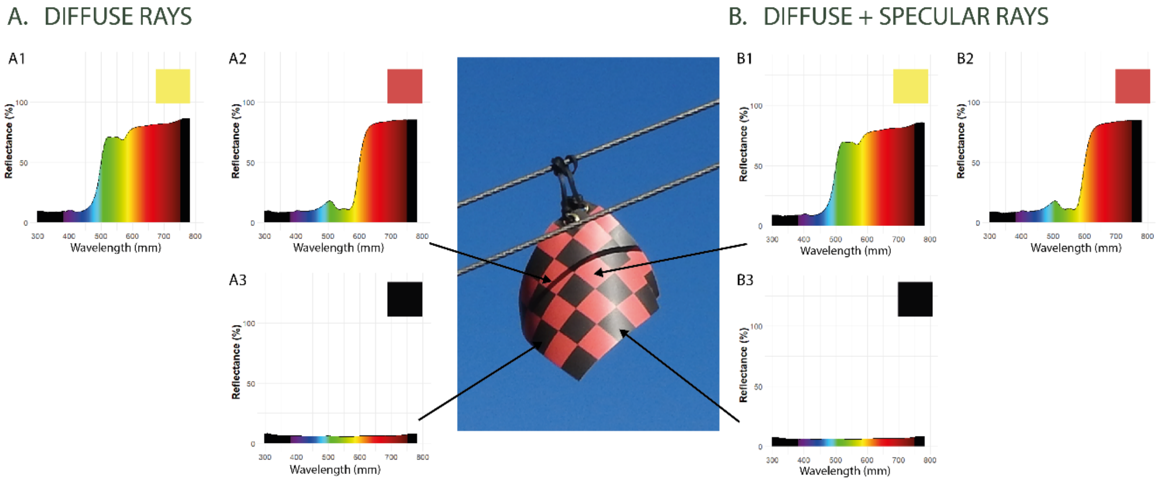


**Figure S3**. **Diffuse and specular reflection of Flags.** A1 and B1 correspond to a yellow square, A2 and B2 correspond to a red square, and A3 and B3 correspond to a black square.

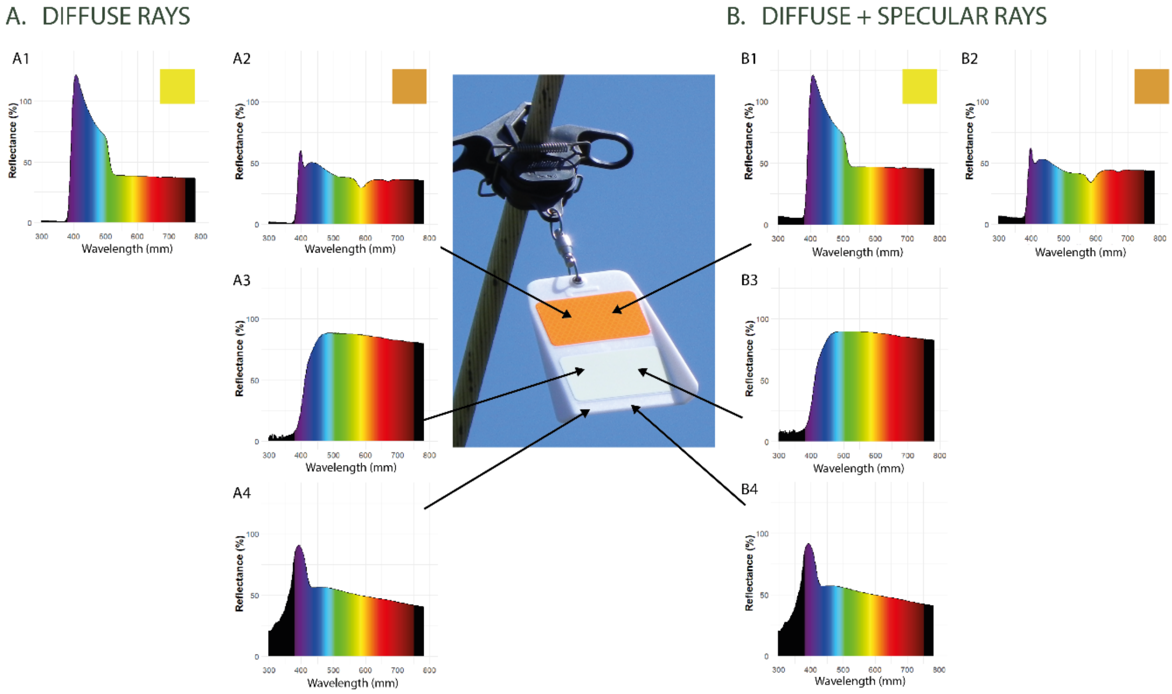


**Figure S4**. **Diffuse and specular reflection of white Fireflies**. A1 and B1 correspond to the yellow retroreflector, A2 and B2 correspond to the orange retroreflector, A3 and B3 correspond to the phosphorescent retroreflector, and A4 and B4 correspond to a white beacon.

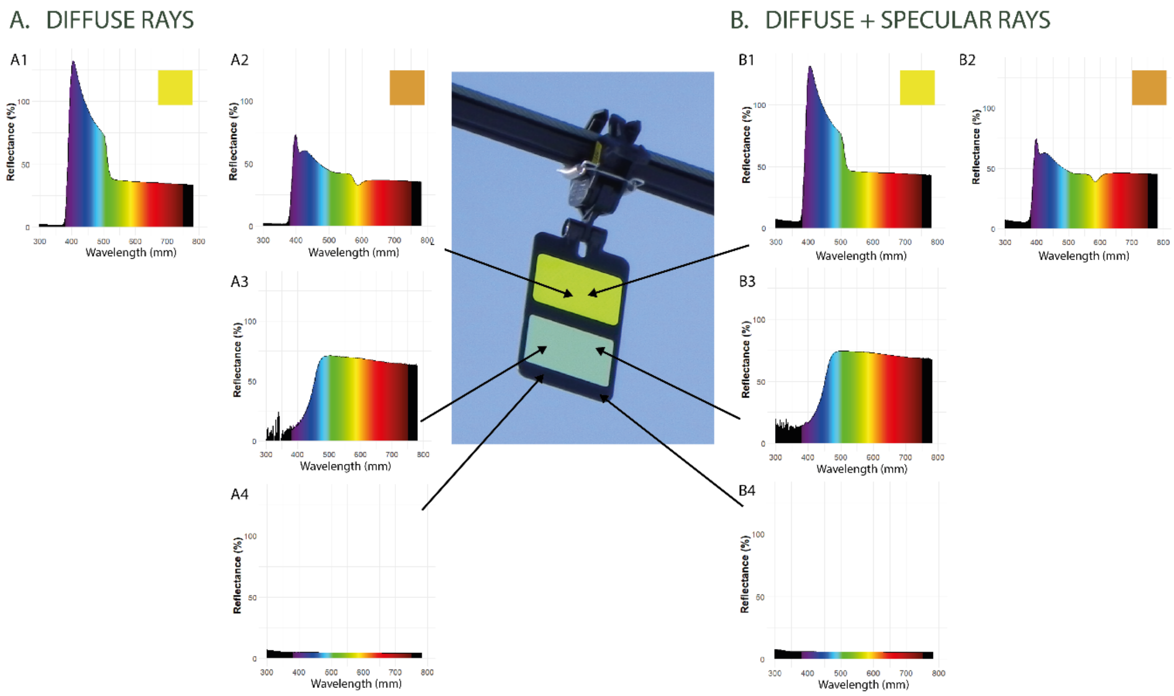


**Figure S5**. **Diffuse and specular reflection of black Fireflies**. A1 and B1 correspond to the yellow retroreflector, A2 and B2 correspond to an orange retroreflector (not shown). A3 and B3 correspond to the phosphorescent retroreflector, and A4 and B4 correspond to a black beacon.

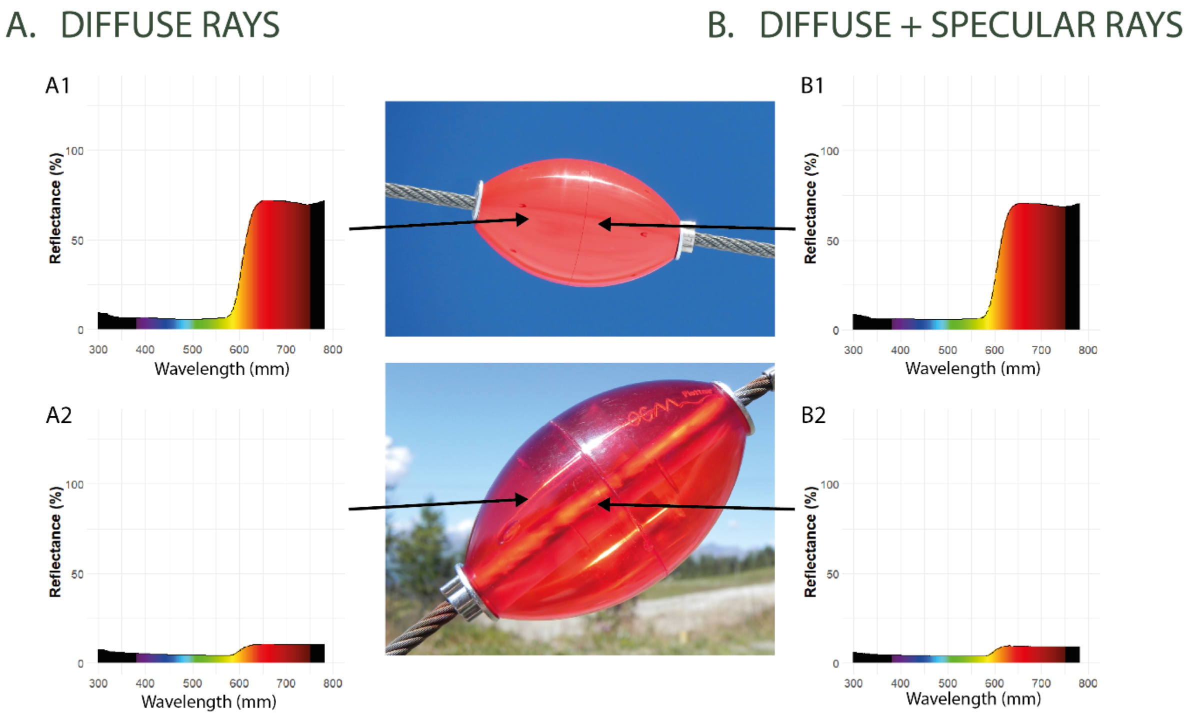


**Figure S6**. **Diffuse and specular reflection of Floats**. A1 and B1 correspond to a transparent float, and A2 and B2 correspond to an opaque float.

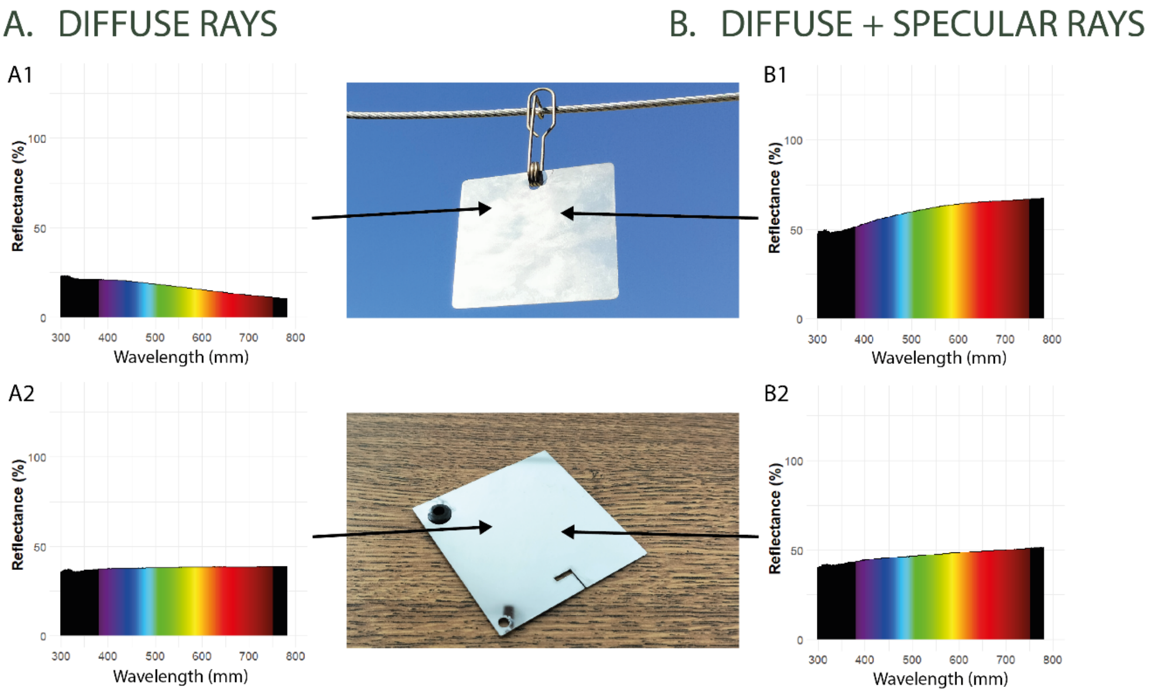


**Figure S7**. **Diffuse and specular reflection of stainless-steel plates**. A1 and B1 correspond to a shiny plate, and A2 and B2 correspond to a matte plate.

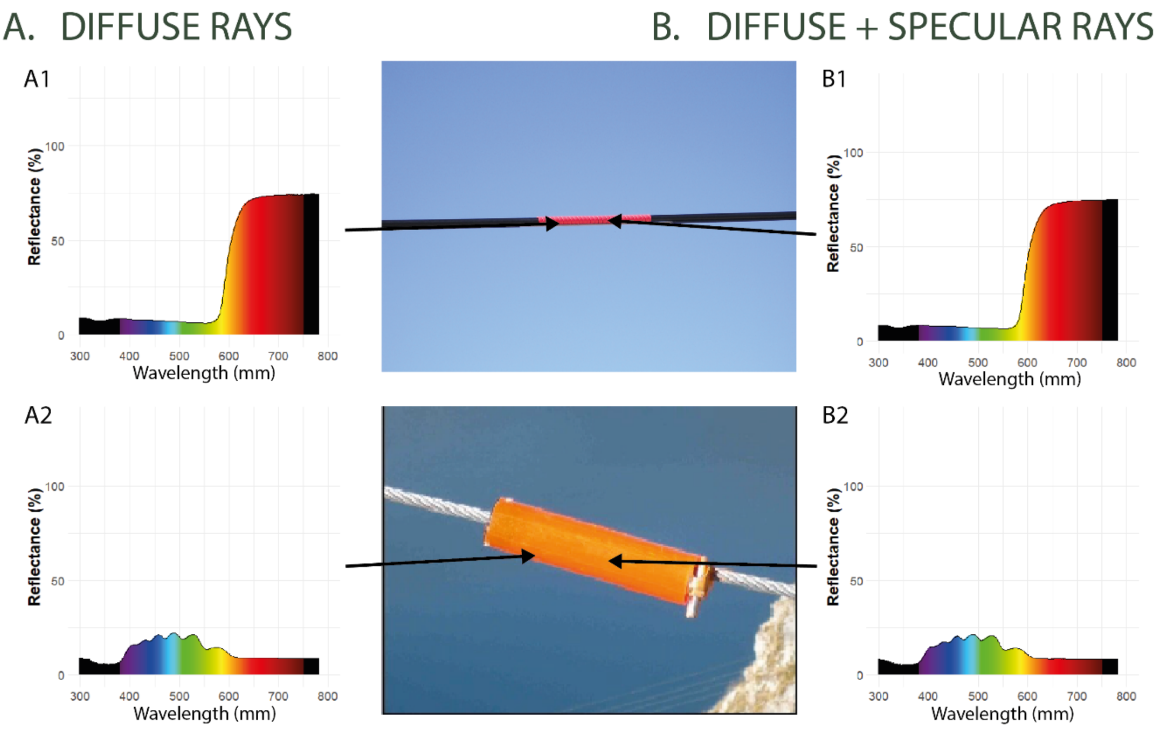


**Figure S8**. **Diffuse and specular reflection of spirals and fluorescent tubes**. A1 and B1 correspond to a red spiral, and A2 and B2 correspond to a fluorescent tube.
